# Crystallographic, kinetic, and calorimetric investigation of PKA interactions with L-type calcium channels and Rad GTPase

**DOI:** 10.1101/2023.10.24.563811

**Authors:** Randy Yoo, Omid Haji-Ghassemi, Jiaming Xu, Ciaran McFarlane, Filip van Petegem

**Affiliations:** Department of Biochemistry and Molecular Biology, University of British Columbia, Life Sciences Institute, 2350 Health Science Mall, Vancouver, British Columbia V6T 1Z3, Canada; Program in Molecular Medicine, the Hospital for Sick Children Research Institute, Toronto, Ontario M5G 0A4, Canada; Department of Biochemistry, University of Toronto, Toronto, Ontario M5G 1M1, Canada; Department of Biological Sciences, University of Calgary 2500 University Drive, N.W. Calgary, Alberta, T2N 1N4, Canada; Evotec, 114 Innovation Drive, Milton Park, Abingdon, OX14 4RZ, UK

**Keywords:** cAMP-depending protein kinase A, voltage gated calcium channel, GTP binding protein, diabetes, synaptic plasticity, cardiac muscle, beta-adrenergic signaling, enzyme kinetics, isothermal titration calorimetry, X-ray crystallography

## Abstract

β-adrenergic signalling leads to activation of cAMP-dependent protein kinase (PKA), which can regulate the activity of L-type voltage-gated calcium channels (Ca_V_s) in multiple tissues. In Ca_V_1.2, various sites have been proposed to be involved, including Ser1981 in the C-terminal tail. Its phosphorylation is linked to diabetes progression, synaptic plasticity, and the augmentation of Ca^2+^ currents in smooth muscle. Its role in augmenting cardiac Ca^2+^ currents has been heavily scrutinized, with alternative models including the sites Ser1718 and Ser1535. Recently, the GTPase Rad has been identified as a critical PKA target that mediates the augmentation of cardiac Ca_V_1.2 currents upon its phosphorylation. However, it is unclear which of the four potential sites (Ser25, Ser38, Ser272, and Ser300) are favored by PKA. Using quantitative binding experiments and enzyme kinetics, we show that there are two Tiers of target sites, with Ca_V_1.2 residue Ser1981 and Rad residues Ser25 and Ser272 forming Tier 1 substrates for PKA. The other sites form a second Tier, with PKA only showing minimal detectable activity. The Tier 1 substrates share a common feature with two arginine residues that anchor the peptide into the active site of PKA. We report crystal structures of the PKA catalytic subunit (PKAc) with and without a Ca_V_1.2 substrate that represent different successive conformations prior to product turnover. Different target sites utilize different anchoring residues, highlighting the plasticity of PKAc to recognize substrates.

**Summary:** Stress signals can alter the electrical properties of excitable cells. cAMP-dependent protein kinase A (PKA) is a key enzyme that is activated upon β-adrenergic stimulation and can alter the function of L-type voltage-gated calcium channels (Ca_V_s) in various tissues. There is a lot of controversy surrounding the exact recognition and specificity of PKA towards Ca_V_1.2, a key calcium channel located in neuronal, cardiac, and smooth muscle tissue, among others. Using a quantitative and unbiased approach, we determined the substrate specificities of PKA towards various sites in Ca_V_1.2 and Rad, an inhibitory protein. Our work highlights two Tiers of substrates, suggesting a potential graded response. Using X-ray crystallography, we determined a high-resolution structure of PKA bound to its strongest target site in Ca_V_1.2, showing how PKA undergoes multiple structural transitions towards binding and how it makes use of a unique anchoring residue.

## Introduction

β-adrenergic signaling can influence electrical excitability through PKA-mediated phosphorylation of ion channels. Particularly relevant is Ca_V_1.2, a voltage gated Ca^2+^ channel that is expressed in multiple tissues, including vascular smooth muscle, neuronal cells, cardiomyocytes, and pancreatic β cells^1^. As Ca^2+^ is a potent intracellular second messenger, fine-tuning the activity of Ca_V_1.2 through phosphorylation plays an important role in various physiological events.

Ca_V_1.2 consists of multiple subunits, including the pore-forming α_1c_, an extracellular α_2_ο and a cytoplasmic β subunit. The α_1c_ subunit consists of four repeats (I-IV) containing six α-helical transmembrane segments each (**Fig. 1A**). The loop connecting repeats I and II (‘I-II loop’) forms a high-affinity binding site for the Ca_V_β subunit^2–4^. With the exception of a small EF-hand domain, the extensive C-terminal tail of the channel has remained invisible in cryo-EM reconstructions of Ca_V_1.2^5,6^, but crystal structures are available for small segments in complex with Calmodulin^7,8^ and Junctophilin^9^ (**Fig. 1A**). Ca_V_1.2 is subject to regulation via many post-translational modifications including phosphorylation^10–12^. Although many kinases have been reported to phosphorylate the channel at various sites, Ser1981 in human Ca_V_1.2 (Ser1928 in rabbit Ca_V_1.2) was suggested to be a major substrate for cAMP-dependent kinase (PKA) and protein kinase C ^13–15^. The phosphorylation of Ser1981 has been suggested to play a role in many important pathophysiological and physiological processes, such as vascular complications in diabetic individuals and in mediating synaptic plasticity in hippocampal neurons through β-adrenergic signalling ^16–18^.

**Figure 1.**
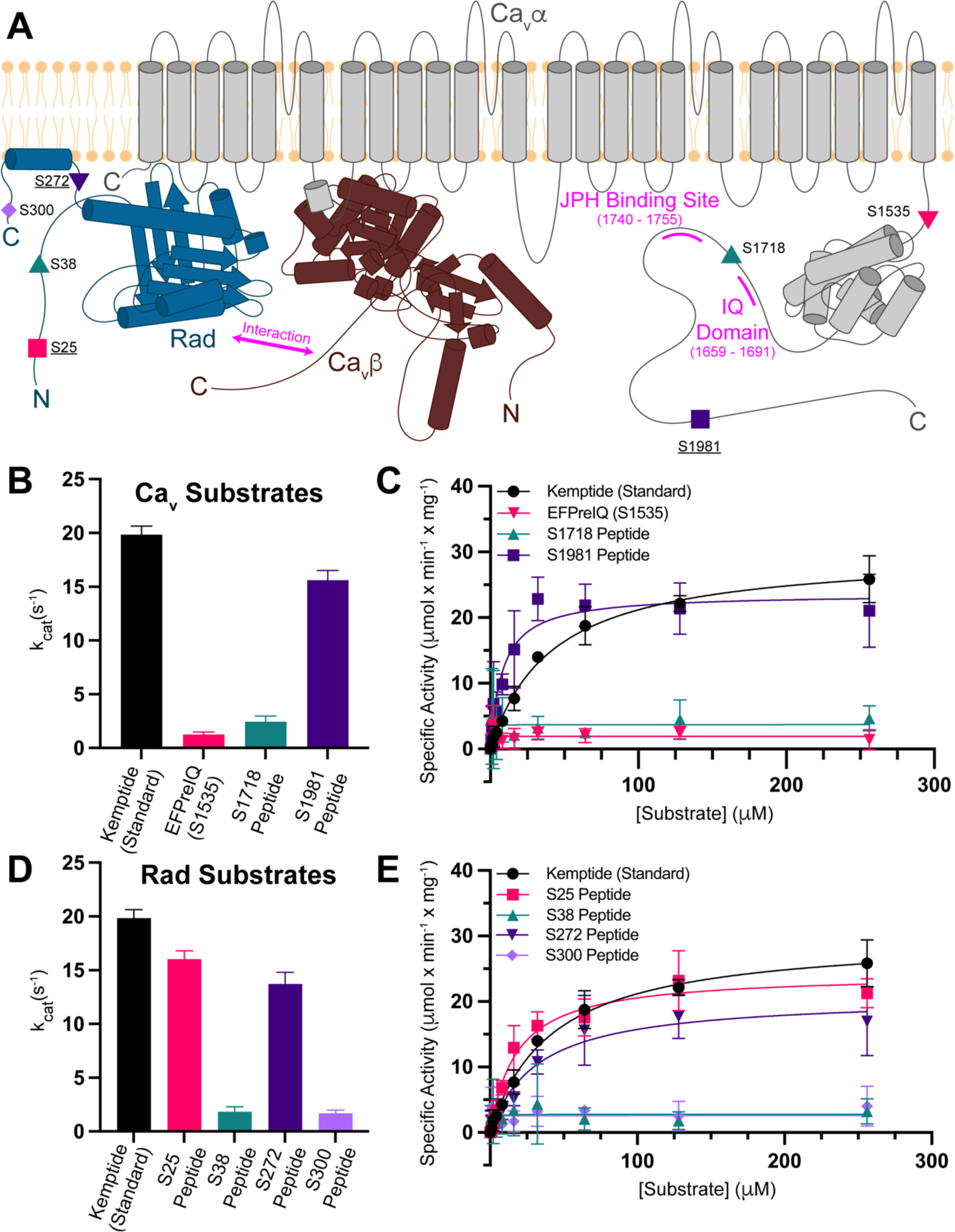
Kinetics of PKAc Phosphorylation of Ca_V_, Rad, and Kemptide Substrates. ***A***, A schematic of the membrane topology of the Ca_V_1.2 channel (α_1c_ subunit in grey and β subunit in brown) and nearby Rad (dark blue) with proposed PKA target sites. The binding sites of junctophilin (JPH) and Calmodulin (IQ domain) are shown in magenta. The Rad C-terminal aliphatic α helix is depicted as being anchored parallel to the plane of the membrane as proposed previously^1^. The underlined sites indicate a PKA site that displayed comparable kinetics to Kemptide. ***B*,** The k_cat_ values determined from the non-linear regressions of kinase assays utilizing: Kemptide (n=3, black), EFPreIQ (Ser1535) (n=3, pink), Ser1718 peptide (n=3, teal), and Ser1981 peptide (n=4, purple) are shown as a bar graph. ***C,*** The average specific activity values of PKAc-mediated phosphorylation measured at various concentrations of the same substrates as in panel B. The solid curves depict the non-linear regression fitted to the data points based on a Michaelis-Menten equation. ***D***, The k_cat_ values determined from non-linear regression fitting of kinase assays utilizing Kemptide (n=3, black) and Rad Peptides: Ser25 peptide (n=3, pink), Ser38 peptide (n=3, teal), Ser272 peptide (n=4, purple), and Ser300 peptide (n=3, violet) are shown as a bar graph. ***E,*** The average specific activity values of PKAc-mediated phosphorylation measured at various concentrations of the same substrates as in panel D. In panels B-D, all error bars represent standard deviations.

Ser1981 phosphorylation has been proposed to cause dissociation of the β_2_ adrenergic receptor (β_2_AR) from the channel upon phosphorylation^15^. It is also thought to augment Ca_V_1.2 currents in various tissues^16,18^, but a clear mechanistic description has been lacking. Studying the effects of Ser1981 phosphorylation is complicated by the fact that in neuronal and cardiac cells, up to 80% of Ca_V_1.2 channels are proteolytically cleaved at Ala1822^19,20^. The resulting distal C terminus (dCT), containing Ser1981, was suggested to serve as both a transcriptional factor and as an autoinhibitory domain^21,22^. The role of Ser1981 in cardiac β-adrenergic signalling has also been questioned^23^. Other phosphorylation sites on Ca_V_1.2 have also been proposed to play a role in channel activation, including Ser1535 just upstream of the EF-hand domain, and Ser1718 located between the channel’s IQ domain and junctophilin-binding motif ^24,25^. Similarly, their importance has also been questioned^26,27^.

Recently, proximity proteomics demonstrated that Rad, a small GTPase that inhibits Ca_V_s, is enriched in the cardiac Ca_V_1.2 micro-environment but is depleted during β-adrenergic stimulation^26^. Activation of PKA in heterologous cells co-expressing both the α_1C_ and β subunits of Ca_V_1.2 with Rad was found to be sufficient in augmenting the Ca_V_1.2 currents^26^, although in a separate study, both Rad-dependent and Rad-independent effects were observed in a heterologous expression system^28^. The model proposes that the PKA-mediated phosphorylation of Rad at four putative residues (Ser25, Ser38, Ser272, and Ser300) causes it to dissociate from the Ca_V_1.2 β subunit, relieving its inhibitory effect and consequently resulting in enhanced peak currents upon excitation of the heart. Mice lacking these sites in Rad show a drastic reduction in β-adrenergic mediated changes in cardiac contractility and in exercise capacity^29^.

With newfound evidence of Rad being the main target of PKA during the fight-or-flight response of the heart, questions remain around the importance of Ser1981, Ser1718 and Ser1535 on Ca_V_1.2. One possibility is that they play roles in different tissues^16,18^. In light of these contradictory reports, we took an unbiased approach, testing how well PKA recognizes each of the three proposed Ca_v_1.2 and four Rad sites (**Table 1**). Using kinase assays and isothermal titration calorimetry experiments (ITC) we determined that Ser1981 on Ca_V_1.2 and both Ser25 and Ser272 on Rad are the most competent substrates of PKA (α isoform). In contrast, residues Ser1535 and Ser1718 on Ca_V_1.2 and Ser38 and Ser300 on Rad were found to be comparatively poorer substrates with no binding detected via ITC. Using X-ray crystallography, we solved structures of the PKA catalytic subunit (PKAc) in complex with the Ca_v_1.2 Ser1981 peptide, which reveals an interaction previously not reported in other PKA:substrate complexes, whereby an aromatic Phe in the P+1 position of the substrate interacts with a hydrophobic pocket of PKA.

**Table 1.**
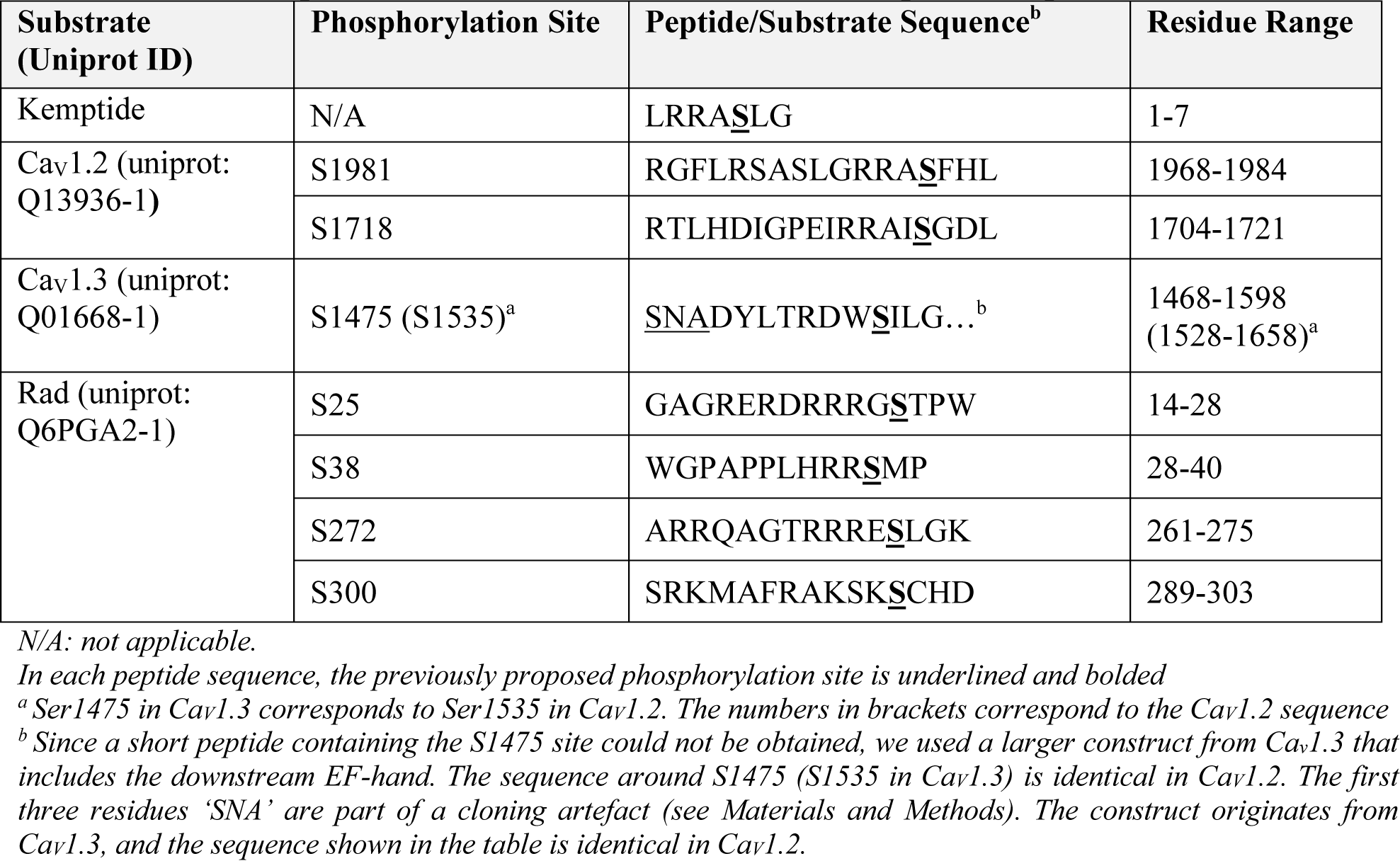
Ca_V_1.2 Peptides, EFPreIQ Construct, and Rad Peptides Sequences.

## Results

To investigate the competency of the proposed PKA target sites, we adapted and performed discontinuous kinase assays using an ADP-GloMax^TM^ Assay kit. The catalytic subunit of PKA (PKAc) undergoes bisubstrate second order kinetics preferring the binding of ATP and Mg^2+^ to occur prior to substrate binding^30–32^. Therefore, we performed assays with excess ATP (1 mM) and Mg^2+^ (5 mM) to saturate PKAc prior to substrate binding. As a positive control, we used Kemptide, a frequently used reference peptide that is readily phosphorylated by PKA (**Table 1**). The kinetic properties we determined are within the range of values previously reported for kemptide, indicating that our assay accurately captures the kinetic properties of PKAc phosphorylation (**Table 2**) (**Fig. 1**)^33–36^. For all residue numbering, we used the sequences for human Ca_V_1.2 (NCBI NP_955630) and mouse Rad (NCBI NP_062636.2). For Rad, mouse was chosen rather than human, as previous experiments suggesting the importance of Rad were carried out with mouse^26^.

**Table 2.** Michaelis-Menten Kinetic Parameters of PKAc with Kemptide, Ca_V_ and Rad Substrates.

| Substrate (sample size) | $k_{\text{cat}}$ ($\text{s}^{-1}$) | $K_m$ ($\mu\text{M}$) | $K_{\text{cat}}/K_m$ ($\text{s}^{-1} \times \mu\text{M}^{-1}$) |
| --- | --- | --- | --- |
| Kemptide (n=3) | $19.8 \pm 0.8$ | $41 \pm 5$ | $0.48 \pm 0.06$ |
| EFPreIQ (S1535) (n=3) | $1.3 \pm 0.2$ | ND | ND |
| S1718 Peptide (n=3) | $2.5 \pm 0.5$ | ND | ND |
| S1981 Peptide (n=4) | $15.6 \pm 0.9$ | $8 \pm 2$ | $1.9 \pm 0.5$ |
| Rad S25 Peptide (n=3) | $16.0 \pm 0.8$ | $18 \pm 3$ | $0.9 \pm 0.2$ |
| Rad S38 Peptide (n=3) | $1.9 \pm 0.4$ | ND | ND |
| Rad S272 Peptide (n=4) | $13 \pm 1$ | $32 \pm 8$ | $0.4 \pm 0.1$ |
| Rad S300 Peptide (n=3) | $1.7 \pm 0.3$ | ND | ND |

*Errors reported as SD.*
*ND: Not determined due to inaccurate $K_m$ as a result of poor fitting.*

### Ser1535 and Ser1718 on Ca_V_1.2 are poor PKA substrates compared to Ser1981

To investigate PKA-mediated phosphorylation at Ser1981 and Ser1718, we used synthetic peptides derived from the human Ca_V_1.2 sequence (**Table 1**). Both Ser1981 and Ser1718 reside in predicted intrinsically disordered regions, and peptides are thus reasonable mimics. The peptides include at least 13 residues upstream of these sites. The Ser1535 site is located only a few residues downstream of the last transmembrane segment. Since a peptide containing this site displayed poor solubility, we instead opted to use a longer construct that also contains the downstream EF-hand domain. The construct corresponding to the Ca_V_1.2 sequence failed to purify, so we instead used the corresponding region in Ca_V_1.3, which could be readily expressed and purified. Importantly, the region around the proposed target site (corresponding to residue Ser1475 in human Ca_V_1.3, **Table 1**) is identical in both isoforms, and the sequence identity in the entire fragment is 96% **(Fig. S1)**. This construct includes only 7 residues upstream of the proposed site, as residues more upstream are part of a transmembrane helix, likely unavailable for binding and their hydrophobic nature would negatively affect solubility.

The Ser1981 peptide is a competent substrate of PKAc, with a k_cat_/K_m_ = 1.9 ± 0.5 s^−1^ × μM^−1^, and is a more robust substrate than the commonly used reference Kemptide (k_cat_/K_m_ = 0.48 ± 0.06 s^−1^ × μM^−1^) (**Table 2**) (**Fig. 1B and 1C**). In contrast, the Ser1718 peptide and Ser1535 mimicking construct are poorer substrates, displaying comparatively lower catalytic activities (**Table 2**) (**Fig. 1B and 1C**). An accurate K_m_ value cannot be calculated as the low catalytic activity at lower substrate concentrations does not allow an accurate fitting. However, the enzyme is easily saturated in these assays with the Ser1718 and Ser1535 substrates displaying a k_cat_ = 2.5 ± 0.5 s^−1^ and k_cat_ = 1.3 ± 0.2 s^−1^ respectively (**Table 2**) (**Fig. 1C**). Therefore, it is possible that Ser1718 and Ser1535 are only significantly phosphorylated by PKA during prolonged β-adrenergic signalling, with Ser1981 representing a more sensitive target.

### Ser25 and Ser272 on Rad are preferentially targeted for PKA-mediated phosphorylation

Due to the pivotal role of Rad in β-adrenergic signalling of the heart, we conducted kinase assays with Rad-derived peptides to determine which sites are preferentially phosphorylated. All four putative PKA sites are in predicted intrinsically disordered regions or flanked by secondary structural elements. Both the Ser25 (k_cat_/K_m_ = 0.9 ± 0.2 s^−1^ × μM^−1^) and Ser272 (k_cat_/K_m_ = 0.4 ± 0.1 s^−1^ × μM^−1^) peptides display kinetic properties comparable to Kemptide (k_cat_/K_m_ = 0.48 ± 0.06 s^−1^ × μM^−1^) (**Table 2**) (**Fig. 1D and 1E**). In contrast, the Ser38 and Ser300 peptides are poor substrates, with k_cat_ values of 1.9 ± 0.4 s^−1^ (Ser38) and 1.7 ± 0.3 s^−1^ (Ser300), but no reliable K_m_ values could be obtained (**Table 2**) (**Fig. 1D and 1E**). Based on these data, Ser25 and Ser272 seem to form the main targets for PKA in Rad.

### PKAc binds with micromolar affinity to Ser1981 of Ca_V_1.2 and Ser25 and Ser272 of Rad

We performed isothermal titration calorimetry (ITC) to determine the thermodynamic properties of PKAc binding to Ca_V_ and Rad substrates in the presence of excess Mg^2+^ (5 mM) and 0.5mM AMP-PNP (a non-hydrolyzable ATP analog) (**Fig. 2**). We could not detect any heat signals indicative of binding when using the Ser1535 mimicking construct (**Fig. 2A**) or with the Ca_V_1.2 Ser1718 peptide (**Fig. 2B**). We only detected binding when titrating the Ca_V_1.2 Ser1981 peptide into PKAc (K_d_ = 34 ± 4 μM) (**Fig. 2C**) (**Table 3**). For the Rad peptides representing phosphorylation sites present on the N-terminal tail of the protein, we detected binding when titrating the Ser25 peptide (K_d_ = 460 ± 20 μM) (**Fig. 2D**) (**Table 3**) but no binding when titrating the Ser38 peptide (**Fig. 2E**) (**Table 3**). For the C-terminal tail phosphorylation site peptides, we detected binding when using the Ser272 peptide (K_d_ = 126 ± 4 μM) (**Fig. 2F**) (**Table 3**), but no binding when using the Ser300 peptide (**Fig. 2G**). The binding of PKAc to the Ca_V_1.2 Ser1981 peptide is endothermic (ΔH = 13 ± 1 kJ × mole^−1^, -TΔS = −36 ± 2 kJ × mole^−1^), in contrast with the exothermic profiles of both Rad Ser25 (ΔH = −10.9 ± 0.3 kJ × mole^−1^, -TΔS = −6.8 ± 0.2 kJ × mole^−1^) and Rad Ser272 (ΔH = −17 ± 1 kJ × mole^−1^, -TΔS = −4 ± 1 kJ × mole^−1^) peptides (**Table 3**). The binding of each peptide is entropically favourable, consistent with the conformational selection model of PKA substrate binding^37^ (**Table 3**). Overall, the ITC data only show significant binding for the same peptides that are more optimal substrates for PKAc.

**Figure 2.**
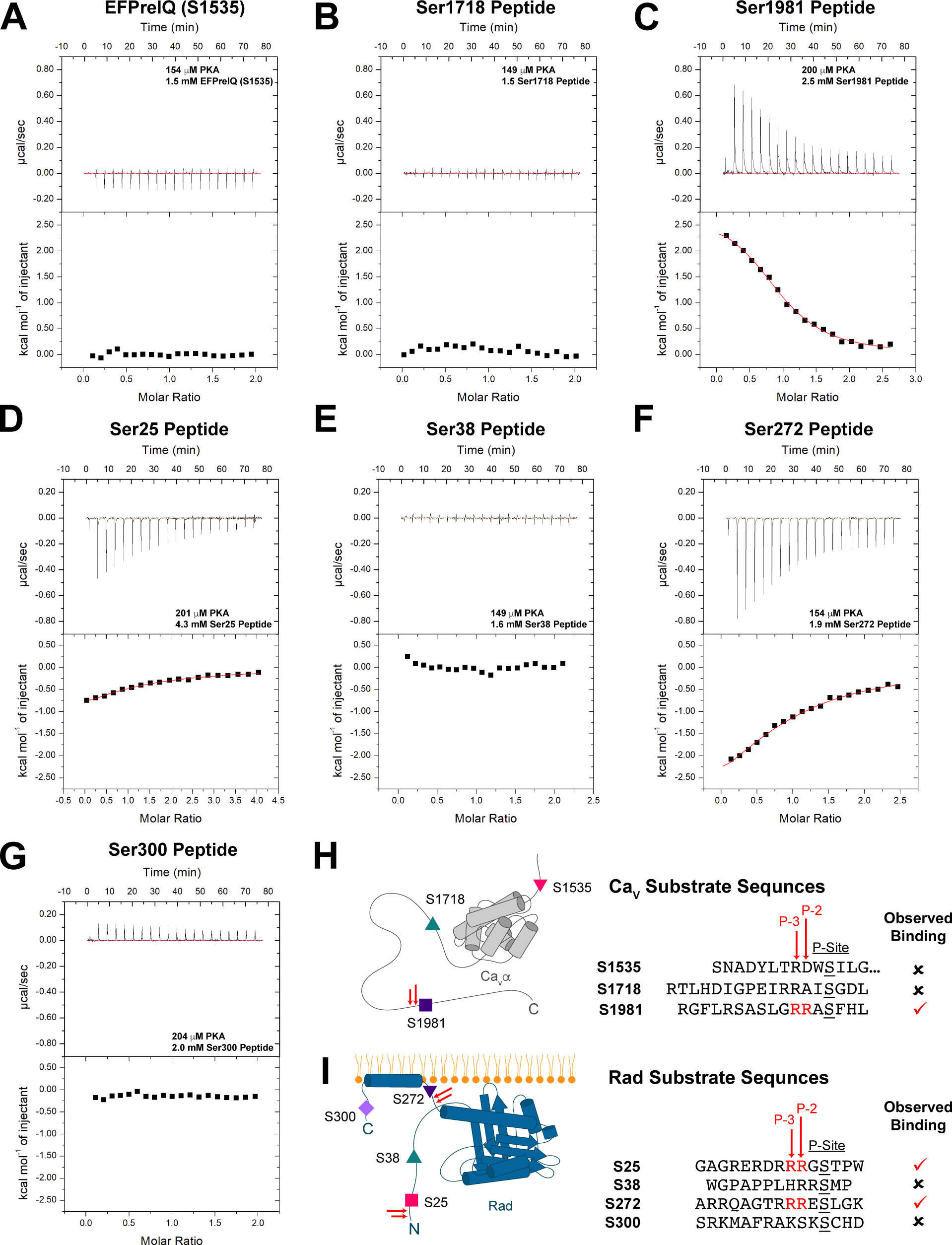
ITC Experiments of PKAc:AMP-PNP Binding to Ca_V_ and Rad Substrates. Representative isotherms from ITC experiments titrating ***A***, 1.5 mM EFPreIQ (S1535); ***B*** 1.5 mM S1718 peptide; ***C***, 2.5 mM Ser1981 peptide; ***D***, 4.3 mM Ser25 peptide; ***E*** 1.6 mM Ser38 peptide; ***F***, 1.9 mM S272 peptide; and ***G***, 2 mM Ser300 peptide into 149-204 μM of PKAc in the presence of 0.5 mM AMP-PNP and 5 mM MgCl_2_. A schematic illustrating a summary of the results of the isothermal titration calorimetry experiments utilizing ***H***, Ca_V_ substrates or ***I***, Rad substrates. Substrates with arginine residues at the P-3 and P-2 positions are indicated using two red arrows.

**Table 3.** Thermodynamic Binding Properties of PKAc:AMP-PNP to Substrates.

| Substrate<br>(replicates) | $\overline{K_D}$ ($\mu\text{M}$) | $\overline{N}$ | $\overline{\Delta G}$<br>($\text{kJ} \times \text{mole}^{-1}$) | $\overline{\Delta H}$<br>($\text{kJ} \times \text{mole}^{-1}$) | $-\overline{T\Delta S}$<br>($\text{kJ} \times \text{mole}^{-1}$) |
| --- | --- | --- | --- | --- | --- |
| Ca<sub>v</sub>1.2 S1981 Peptide (n=3) | $34 \pm 4$ | $0.98 \pm 0.05$ | $-23.7 \pm 0.2$ | $13 \pm 1$ | $-36 \pm 2$ |
| Ca<sub>v</sub>1.2 S1718 Peptide (n=1) | ND | ND | ND | ND | ND |
| Ca<sub>v</sub>1.3 EFPreIQ (S1535) (n=1) | ND | ND | ND | ND | ND |
| Rad S25 Peptide (n=3) | $460 \pm 20$ | $0.83 \pm 0.06$ | $-17.7 \pm 0.1$ | $-10.9 \pm 0.3$ | $-6.8 \pm 0.2$ |
| Rad S38 Peptide (n=2) | ND | ND | ND | ND | ND |
| Rad S272 Peptide (n=3) | $126 \pm 4$ | $0.98 \pm 0.05$ | $-20.70 \pm 0.08$ | $-17 \pm 1$ | $-4 \pm 1$ |
| Rad S300 Peptide (n=1) | ND | ND | ND | ND | ND |

Peptides were titrated into the cell containing PKAc. Measurements were performed at 4 °C in 50mM HEPES (pH 7.4), 150mM KCl, 5mM MgCl<sub>2</sub>, and 2mM TCEP. Errors reported as standard deviations. ND: Not determined due to unobservable binding or poor fit.

### Structures of ApoPKAc, AMP-PNP PKAc, and PKAc-Ser1981 ternary complexes

Given our findings that the Ca_V_1.2 Ser1981 peptide, Rad Ser25 peptide, and Rad Ser272 peptides are all good PKA substrates, we attempted to capture PKAc ternary complexes with each of these substrates to help better understand PKA substrate specificity. Our efforts yielded two crystal structures of PKAc ternary complexes (Complex1 and Complex2) of PKA bound to both AMP-PNP/Mg^2+^ and the Ca_V_1.2 Ser1981 peptide at 2.85Å and 2.99Å resolutions, respectively **(Table S1)**. Thus far, complexes with the Rad peptides escaped crystallization, likely a result of their weaker affinity for PKAc. We also collected a dataset for PKAc in the presence of the Ser1718 peptide. This 2.75Å data set contains four PKAc molecules in the asymmetric unit referred to here as Apo/AMP-PNP PKAc, as it lacks density for the Ser1718 peptide. Three out of the four molecules in the asymmetric unit are in the apo state (Chain C: ApoPKAc1, Chain F: ApoPKAc2, Chain H: ApoPKAc3) and one is bound to AMP-PNP (Chain D). Altogether, our structures depict PKAc in different catalytic stages and conformations.

### Dynamic nature of the PKAc catalytic cycle

The PKAc conformations are defined as being in the open, intermediate, or closed states. This classification is based on the distances of the alpha carbons of Gly52 and Asp166, the C_α_ atoms of Ser53 and Gly186, the Nε2 atom of His87 and closest oxygen in pThr197 side chain, and the backbone oxygen of Glu170 and the hydroxyl group of Tyr330^33^. Based on these criteria, our Apo structures (apoPKAc) depict an open state of the kinase, whereas the AMP-PNP and ternary complexes correspond to different intermediate states (**Fig. 3A-3D**). The distances between these residue pairs decrease in length in the order: ApoPKAc3 > ApoPKAc1 > ApoPKAc2 > AMP-PNP PKAc > Complex 2 > Complex 1 **(Table S2)**. Thus, these structures represent various states of the kinase starting from the open/unbound state to a fully bound, intermediate state prior to subsequent substrate turnover.

**Figure 3.**
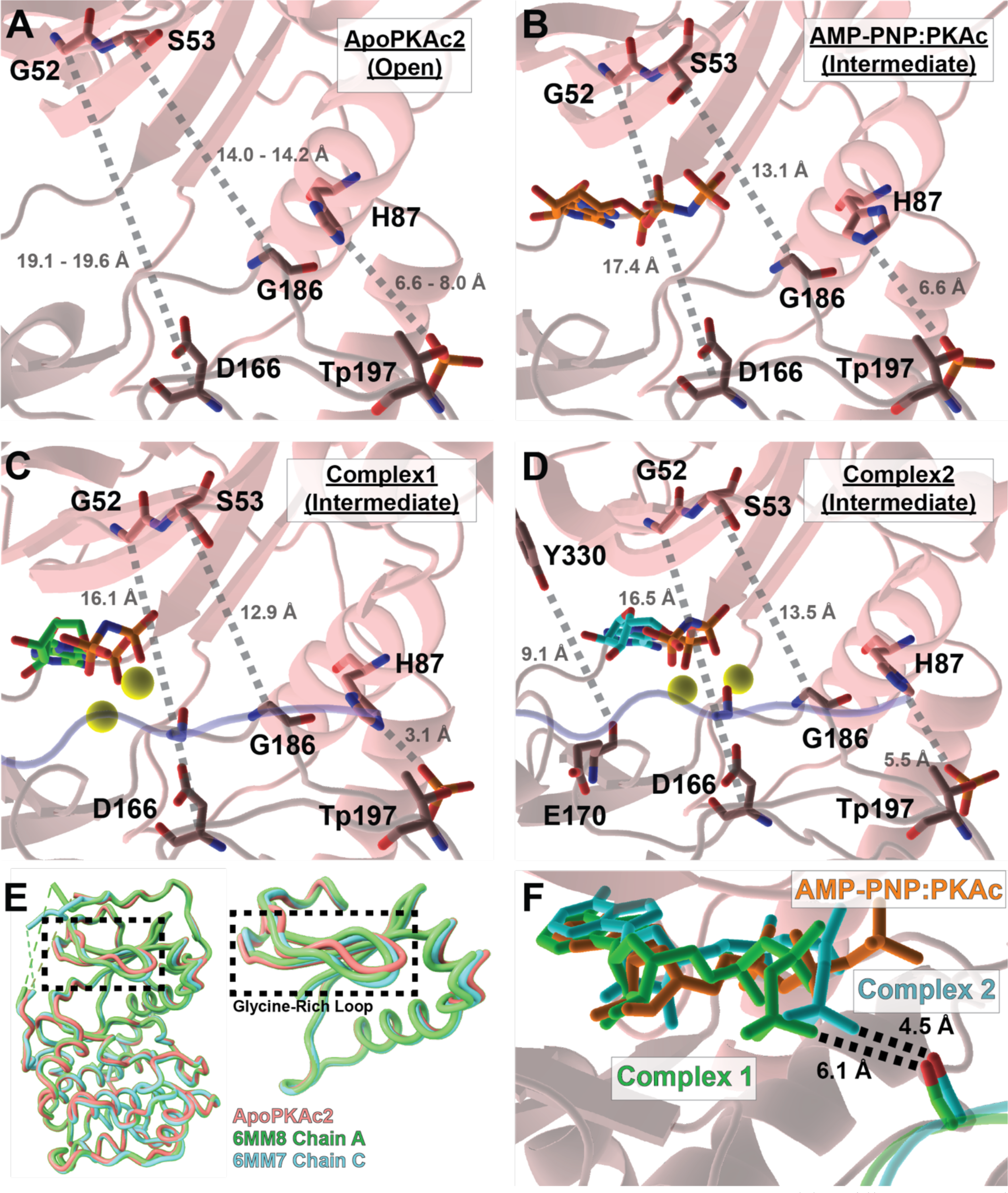
Conformational Differences of the PKAc Structures. ***A-D***, Residues/distances used to classify the open/intermediate/closed state of PKAc molecules are shown. The distances listed for ApoPKAc2 are listed as the range of distances observed in all three ApoPKAc molecules (ApoPKAc1, ApoPKAc2, and ApoPKAc3). The PKAc molecules are depicted as cartoons and sticks. The sticks are coloured according to the following scheme: oxygen atoms, red; nitrogen atoms, blue; and phosphorus atoms, orange. The cartoon and carbon atom sticks are coloured either salmon (small lobe) or dark salmon (large lobe). The Ca_V_ Ser1981 peptide is coloured purple as a ribbon representation and the Ser1981 side chain is shown as sticks. Magnesium ions are depicted as yellow spheres. The AMP-PNP molecules are displayed as sticks with carbon atoms coloured *B,* orange; *C,* green; and *D,* cyan. ***E***, Two PKAc structures from RyR2 ternary structures with the lowest RMSDs and highest similarity score (Z-score) calculated via the DALI server ^2^ are depicted as cartoons and are overlaid the ApoPKAc2 structure (PDB: 8UKN, Chain F). The glycine-rich loop is highlighted using a dashed black box. ***F***, Differences in the substrates of AMP-PNP:PKAc, Complex1, and Complex2 molecules are shown. The PKAc molecule is shown as ribbons and coloured according to the previous scheme. The substrates (AMP-PNP and Ca_V_ Ser1981 peptide) are coloured orange (AMP-PNP:PKAc), green (Complex1, PDB: 8UKP), and cyan (Complex2, PDB: 8UKO). The AMP-PNP molecules are shown as sticks. The Ser1981 side chain is shown as sticks with its hydroxyl group coloured red.

Most notably the ApoPKAc2 molecule adopts a conformation distinct from the other two ApoPKAc molecules in the asymmetric unit (**Fig. 3E**). A DALI search^38^ of ApoPKAc2 yields top hits of some of our previously published PKAc:RyR2 ternary structures with RMSD values of 0.6-0.9 Å **(Fig. S2)**. The main difference observed between these structures and ApoPKAc2 is the glycine-rich loop (residues 47-56) (**Fig. 3E**). This is expected, as the stable positioning of the loop is largely dependent on the presence of a bound nucleotide and can vary greatly in ApoPKAc structures^39^. Nevertheless, the ApoPKAc2 molecule adopts a conformation that most closely resembles a substrate-bound state. This lends support to the conformational selection model as this conformation of PKAc appears to be thermodynamically stable and can be sampled both in the presence and absence of substrate^37,40,41^. The dynamic nature of these structures extends to the active site, as the gamma phosphate of the AMP-PNP molecule adopts various positions in our structures (**Fig. 3F**). Interestingly, despite Complex1 representing a more closed state than Complex2, in the latter the gamma phosphate is more readily primed for phosphoryl transfer, indicated by its shorter distance from the Ser1981 hydroxyl group (**Fig. 3F**). Altogether these observations highlight the dynamic nature of the enzyme both in the presence and absence of substrate.

### PKAc specificity to Ser1981

Our PKAc:substrate ternary complexes contain strong density for residues Gly1969 – Phe1982 of the Ser1981 peptide **(Figs. S3 and 4A).** As evident from our kinase assays and ITC experiments, the two arginine residues at the P-2 and P-3 positions are critical for PKA substrate specificity, as only the peptides containing these showed detectable binding. These residues have previously been found to be important for interactions with both substrates and inhibitory peptides^33,37,42,43^. The arginines mediate multiple interactions with PKAc (**Fig. 4B**). The P-3 Arg side chain participates in a salt bridge interaction with PKAc Glu128. The P-2 Arg side chain fits into an electronegative pocket of the active site formed by the side chains of Glu170, Glu203, and Glu230, forming salt bridges with all three. Outside of these arginine residues, the Ser1981 peptide engages in multiple hydrogen bonding interactions. The Arg133 side chain of PKAc hydrogen bonds with the backbone oxygen of the P-5 Leu1976 (**Fig. 4C**). In addition to its side chain, the backbone oxygen of the P-2 Arg1979 also forms a hydrogen bond with the side chain of Lys168 of PKAc (**Fig. 4D**). Finally, the P+1 Phe1981 nitrogen backbone atom forms a hydrogen bond with the Gly200 oxygen backbone atom (**Fig. 4E**). In addition to these interactions, both PKAc:AMP-PNP:Ca_V_1.2 Ser1981 peptide complexes present an interaction mode not yet seen in PKAc:substrate complexes. The P+1 phenylalanine side chain fits into a hydrophobic pocket of the enzyme, formed by the side chains of PKAc Leu198, Pro202, and Leu205 (**Fig. 4F**). Finally, extensive vdW interactions are observed through the N-terminal portion of the peptide (**Fig. 4A**) involving Phe1970 (P-4), Leu1971(P-5), Arg1972 (P-6), Ser1975 (P-9), Leu1976 (P-10), and Gly1977 (P-11) with various PKAc residues **(Table S3 and Fig. S4)** with a total buried surface area of 257.2 Å^2^ in Complex1 and 210.21 Å^2^ in Complex2.

**Figure 4.**
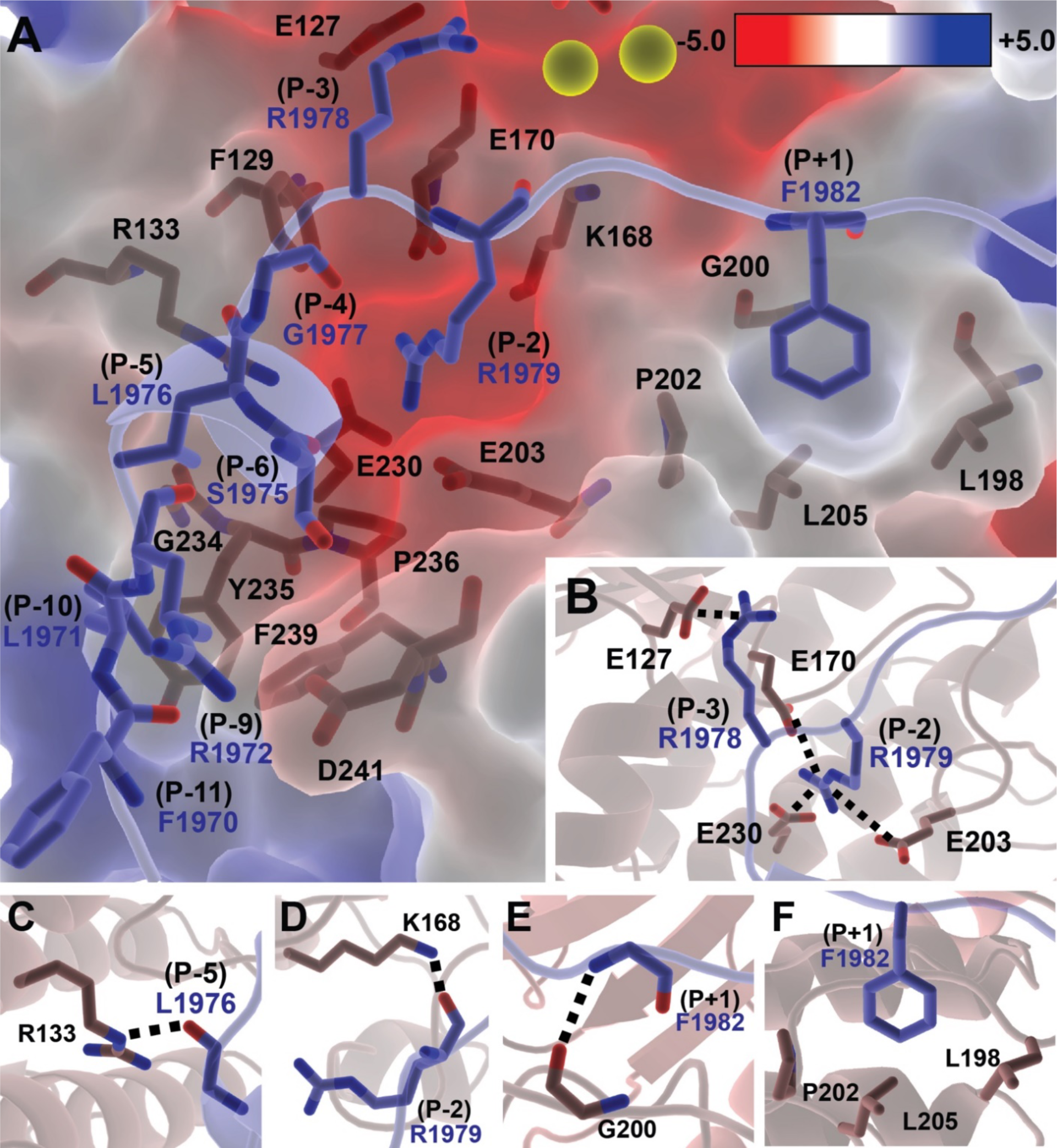
Critical Binding Determinants for PKAc Specificity to the Ca_V_1.2 Ser1981 Peptide. Residues involved in enzyme-peptide recognition are shown as sticks. The Ca_V_1.2 peptide is shown in purple and PKAc in salmon/dark salmon (small lobe/large lobe). Nitrogen atoms are coloured in red and oxygen in blue. ***A***, A surface representation of the active site of PKAc is coloured according to electrostatic potential. Residues involved in vdW and electrostatic interactions are shown as sticks. ***B***, An electronegative pocket is formed by E170, E203, and E230 in which the R1979 side chain at the P-2 position of the S1981 peptide slots into forming salt bridging interactions with E230, E170, and E203. The R1978 in the P-3 position forms a salt bridge with E127. ***C***, The P-5 leucine of the S1981 peptide forms a hydrogen bond with R133 through its backbone oxygen atom (dashed line). ***D***, The mainchain oxygen of R1979 forms a hydrogen bond with the K168 side chain (dashed line). ***E***, The oxygen backbone atom of G200 forms a hydrogen bond with the backbone nitrogen atom of F1982 (dashed line). **F**, A hydrophobic pocket formed by L198, P202, and L205 is occupied by the F1982 side chain of the peptide forming extensive hydrophobic contacts.

Although many structures of PKAc have been reported, only two other ternary PKAc:substrate complexes have been reported. Figure 5 compares the binding of Ca_V_1.2 (Ser1981 peptide) (**Fig. 5A**) with RyR2 (**Fig. 5B**) and phospolamban (PLN) (**Fig. 5C and 5D**), highlighting the divergence in the interactions. In all structures, the P-2 and P-3 arginine side chains are at near identical positions (**Fig. 5E**). The backbone nitrogen atom of the P+1 atom of each substrate engages in a hydrogen bond with Gly200 backbone oxygen atom of PKAc (**Fig. 5F**). Although all substrates engage in extensive vdW interactions at the N-terminal portion of the substrate, they utilize different residues **(Fig. S4)**. The P-5 backbone hydrogen bonding interaction with Arg133 observed in Ca_V_1.2 is also present in RyR2 but not in PLN (**Fig. 5G**). The Ca_V_1.2 Ser1981 peptide differs from both PLN and RyR2 in the usage of a P-7 residue (**Fig. 5H**). In Ca_V_1.2, this residue is an alanine that does not form specific interactions, in contrast with RyR2 where a Tyr residue packs into a shallow hydrophobic groove of PKAc and forms a hydrogen bond with Glu203. In PLN (PDB: 3O7L), an Arg at this position either forms a salt bridge with Glu203 (PDB: 3O7L) or forms a hydrogen bond with the backbone oxygen atom of Arg134 (PDB: 7E0Z) **(Fig. S5)**. Thus, the precise interactions differ in both the N-terminal and C-terminal regions of the substrate peptide.

**Figure 5.**
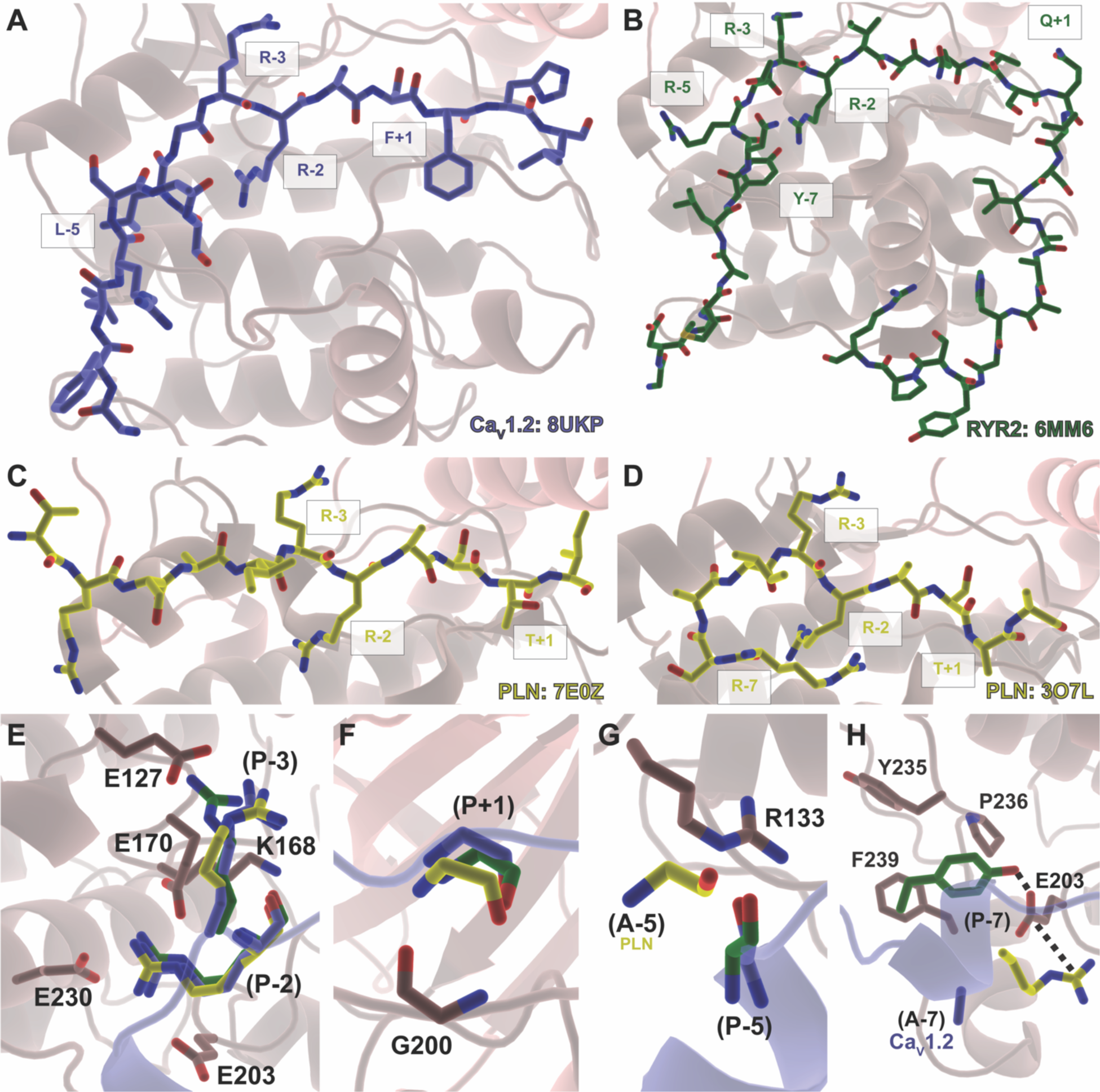
Comparison of PKAc substrate interactions to CaV1.2. Structures of PKAc in complex with ***A***, Ca_V_1.2 Ser1981 peptide (purple) (PDB: 8UKP) ***B***, RyR2 phosphorylation domain (green) (PDB: 6MM6), and ***C*** *and **D**,* Phospholamban (yellow) (PDB: 7E0Z and 3O7L respectively) with substrates depicted as sticks. Residues important for substrate recognition and specific are labelled. ***E***, P-2 and P-3 arginine residues are present in all structures and positioned similarly in all substrates and interacts through various salt bridges and hydrogen bonds. ***F***, The backbone atoms of G200 of PKAc interacts with the P+1 backbone in all PKAc ternary structures. ***G***, RyR2 and Ca_V_1.2 interact with R133 through the backbone oxygen of the P-5 residue. The equivalent residue in PLN does not. ***H***, RyR2 and PLN (PDB: 3O7L but not 7E0Z) P-7 residue side chain interacts through a hydrogen bond and salt bridge respectively to E203. Ca_V_1.2 harbours an alanine at this position and thus lacks a hydrogen bond donor/acceptor to engage in similar interactions.

## DISCUSSION

The results of our kinase and ITC experiments indicate that Ser1981 of Ca_V_1.2 and Ser25 and Ser272 of Rad are most readily phosphorylated by the catalytic subunit of the PKA α isoform (**Fig. 1**). Despite numerous reports describing the importance of Arginines at the P-2 and P-3 positions^44^, several sites lacking these pairs have been proposed as robust PKA targets in Ca_V_1.2 and Rad. Our data underscore the importance of these Arg residues for substrate recognition, as only the peptides containing these showed significant binding and kinase activity (**Fig. 1 and 2**) (**Table 2 and 3**). However, we cannot rule out that the other investigated sites are not being phosphorylated *in vivo,* as our kinase assays do indicate some enzymatic activity, albeit very low. Other factors may also contribute to significant phosphorylation in physiological settings, such as the presence of A-kinase anchoring proteins which have been proposed to associate with Ca_V_1.2, thus enhancing the local concentration of substrate sites next to PKA^45–49^. Thus, even poor substrates can become significantly phosphorylated *in vivo.* However, this local concentration would also apply to the robust target sites, suggesting the presence of two Tiers of substrates. This opens the door for a graded response, whereby short-term β-adrenergic stimulation results in phosphorylation of primarily Ser1981 on Ca_V_1.2 and Ser25+Ser272 on Rad. Longer, more chronic stimulation, perhaps together with reduced activity of phosphatases, may lead to significant phosphorylation of the additional sites. Such a graded response may also have functional consequences not captured with short-term β-adrenergic stimulation.

The C-terminal tail of Rad contains a polybasic α-helix that is thought to mediate interactions with the plasma membrane, enabling the ability of Rad to inhibit Ca_V_1.2. Ser272 resides in this helix, and its phosphorylation thus adds a negative charge that may reduce the affinity for the plasma membrane, resulting in a disinhibition of the channel^50^. However, the involvement of the residues in fully disordered regions should not be ignored. It has been previously observed that phosphorylation of intrinsically disordered regions can affect neighboring structured domains^51–53^. In the case of RyR2, for example, a phosphomimetic at site Ser2814 was shown to induce α-helicity, affecting the affinity for PKA^33^. Thus, the possibility still exists that phosphorylation of other residues in Rad imparts structural changes.

Our study also provides additional insights into substrate recognition by PKAc (**Fig. 5**). Although the binding of PKA to inhibitory peptides is well documented, structures with physiological substrates have been scarce, likely due to their intrinsically weak affinity for PKA. With the current study included, there are now three enzyme:substrate complexes available for PKAc: 1) the cardiac RyR2 phosphorylation hot spot domain containing Ser2808^33^, 2) PLN^37,42^, and 3) the Ca_V_1.2 C-terminal tail containing Ser1981 (this study). Although some interactions appear conserved among the substrates, there are several distinct differences. For example, an aromatic residue at P-7 in RyR2 is buried into a shallow hydrophobic pocket in PKAc. In contrast, PLN utilizes a salt bridge at this position. In Ca_V_1.2 (Ser1981 site), the P-7 position does not contribute, but instead an aromatic anchor at the P+1 position provides additional stabilization. Of note, kinase:substrate interactions are typically weak, but the PKA inhibitor PKI binds to PKAc with high affinity^43^. Interestingly, PKI makes use of both the P-7 and P+1 positions, which contributes to it much higher affinity **(Fig. S6)**.

To our knowledge, no physiological PKAc substrate displays higher than mid-micromolar affinity. This relatively poor affinity of PKAc for its substrates is likely important for the catalytic turnover. It has been proposed that tight binding of a substrate could limit turnover rates by prolonging product release^37^. This is supported by data in which kinetic assays utilizing a phosphorylation-competent PKI peptide, which binds at nM affinity, appears to be a relatively poor substrate when compared head-to-head to Kemptide^43^. Additionally, despite being phosphorylation-competent, the RII subunit binds to PKAc with nM affinity^54,55^ and exhibits “single turnover autophosphorylation” suggested by the non-detectable kinetic activity of the kinase^56,57^. Altogether, these data support the notion that PKAc substrates must balance specificity and affinity to mediate adequate levels of turnover.

Mutations in Ca_V_1.2 have been linked to several disorders, including Brugada, Long QT and Timothy syndromes^58–60^. Mining of the clinvar database (https://www.ncbi.nlm.nih.gov/clinvar/) shows that several sequence variants, found in patients with Long-QT, affect residues in the Ser1981 peptide (**Table S4**). This includes variants in the target serine itself (S1981P, S1981F), as well as the two critical P-2 and P-3 Arg residues (R1978Q, R1979K). Such variants undoubtedly strongly diminish phosphorylation by PKA, through removal of critical contacts. Other variants likely introduce steric hindrance (G1969A, G1977S, G1977D) or restrict the backbone flexibility (S1973P, A1974P). Although the cause of pathogenicity remains to be tested, the clustering of several variants found in Long-QT patients suggests a potential role for phosphorylation of Ser1981 in normal cardiac function, even if it’s not the primary site responsible for Ca_V_1.2 augmentation in cardiac myocytes.

Our work provides insight into the PKA-mediated phosphorylation of Rad elucidating that both Ser25 and Ser272 are phosphorylated by PKA more readily than Ser38 and Ser300. We conclude that the presence of P-2 and P-3 Arginines are critical for PKA specificity whereas other aspects such as backbone atom hydrogen bonding interactions via the P-2 and P+1 with Lys168 and Gly200 respectively are important as they are observed in all PKAc:substrate structures published to date. Finally, different substrates utilize interactions that are not observed in other substrates exemplifying the plasticity in which PKAc can recognize its substrate. Although we were unsuccessful in co-crystallizing the Rad peptides with PKA, further insights may be obtained from such structures, and it will be of interest to test whether the phosphorylation of these peptides results in changes in secondary structure.

## MATERIALS AND METHODS

### DNA Constructs

A mouse PKAcα (cAMP-dependent kinase catalytic subunit, isoform alpha) (UniProtKB/Swiss-Prot: P05132-1) DNA sequence and a human Ca_V_1.3 DNA sequence were cloned into a modified pET28 vector containing an N-terminal 6xHis tag, maltose-binding protein (MBP) tag, and tobacco etch virus (TEV) protease cleavage site. The cloned PKAcα sequence consisted of residues 16-351 (pET28HMT-PKAcα_16-351_). The Ca_V_1.3 construct consisted of residues 1468-1598 (pET28HMT-EFPreIQ) (UniProtKB/Swiss-Prot: Q01668-1) which make up the EF-Hand and PreIQ domain of Ca_V_1.3.

### Recombinant PKAc Protein Expression and Purification

The pET28HMT-PKAcα_16-351_ plasmid was transformed into *Escherichia coli* Rosetta (DE3)pLysS competent cells and plated against chloramphenicol (34μg/ml) and kanamycin (50μg/ml) selection for 16 hours at 37°C. Flasks containing autoinduction media (57) were inoculated and grown at 37°C for 6 hours at 180 RPM before changing the temperature to 18°C. The cells were then grown for an additional 64 hours before harvesting in which cell pellets were stored at −20°C.

All steps in lysis and purification were performed at 4°C or on ice. Upon thawing, cell pellets were lysed via sonication in a lysis buffer containing 20 mM 4-(2-hydroxyethyl)-1-piperazineethanesulfonic acid (HEPES) (pH 7.4), 250 mM KCl, 10 mM imidazole, 10% glycerol, 10 mM MgCl_2_, 1 mM phenylmethylsulfonyl fluoride (PMSF), 1 mM tris(2-carboxyethyl)phosphine hydrochloride (TCEP), 25 μg/mL DNaseI, and 25 μg/mL lysozyme. The lysed cell contents were pelleted via centrifugation (Beckman Coulter, Avanti J-E) at 40,000 × g for 35 min. The soluble lysate was filtered with a 0.45 µm nylon syringe filter, and then applied to a HisTrap Fast Flow IMAC column (GE Healthcare Lifesciences) pre-equilibrated with buffer A (20 mM HEPES [pH 7.4], 250 mM KCl). The column was washed with 15 column volumes (CV) of buffer A before eluting using buffer A + 400mM imidazole. The elution was pooled, spiked with 2mM EDTA, 2mg of TEV and dialyzed in dialysis buffer (20mM HEPES, 250mM KCl, and 10mM β-mercaptoethanol [β-ME]) overnight. The dialyzed sample was then applied to an Amylose Column (New England Biolabs) pre-equilibrated with buffer A. The flowthrough was then collected and applied to a PorosMC column (ThermoFisher Scientific) pre-equilibrated with buffer A. The flowthrough was collected and dialyzed against 10 mM KH_2_PO_4_ (pH 6.6), 10 mM KCl, and 10 mM β-ME for 3 hours. The dialyzed sample was applied to a ResourceS (GE Healthcare) column pre-equilibrated with 10 mM MES (pH 6.6), 10 mM KCl, and 15 mM β-ME and eluted over a 0% to 23% gradient of 10 mM MES (pH 6.6), 1 mM KCl, and 15 mM β-ME over 25 CV. Fractions containing recombinant PKAc verified via SDS-PAGE were pooled and concentrated to 250μL using an Amicon concentrator (10K MWCO; Millipore). The concentrated sample was then applied to a preparative-grade Superdex200 column (GE Healthcare) pre-equilibrated with ITC/Assay buffer (50mM HEPES [pH 7.4], 150mM KCl, 5mM MgCl_2_, and 2mM TCEP). Fractions containing soluble and monomeric PKAc were then pooled and concentrated to 140-300μM for use in ITC experiments or concentrated to 500μM, spiked with 30% glycerol, flash-frozen, and stored at −80°C for later use in kinase assays.

### Recombinant Ca_V_1.3 EFPreIQ Protein Expression and Purification

The pET28HMT-EFPreIQ plasmid was transformed into *Escherichia coli* Rosetta (DE3)pLysS competent cells and plated against chloramphenicol and kanamycin selection for 16 hours at 37°C. Colonies were then inoculated into a 100 mL 2YT media (16 g/L Tryptone, 10 g/L yeast extract, 5 g/L NaCl) starter culture and grown for 16 hours at 37°C. 10 mL of starter culture was then inoculated into 1 L of 2YT media and grown at 37°C at 180 RPM until an OD_600_ of 1.0 was reached. Cultures were then allowed to cool to 18°C and induced to express with 0.4 mM IPTG for 16 hours. The cells were then harvested and stored at −20°C.

All steps in lysis and purification were performed at 4°C or on ice. Upon thawing, cell pellets were lysed via sonication in a lysis buffer (20 mM HEPES [pH 7.4], 500 mM NaCl, 0.2% (v/v) EDTA-free Protease Inhibitor Cocktail III (Millipore), 25 μg/ml DNaseI, and 50 μg/ml lysozyme). The lysed cells were pelleted via centrifugation (Beckman Coulter, Avanti J-E) at 40,000 × g for 35 min. The soluble lysate was filtered with a 0.45 µm nylon syringe filter, and then applied to HisTrap Fast Flow IMAC column (GE Healthcare) pre-equilibrated with buffer B (20 mM HEPES [pH 7.4], 500 mM NaCl). The column was washed with 15 CV of buffer B before eluting the recombinant protein using buffer B + 250 mM imidazole. The elution was pooled and applied to an Amylose Column (New England Biolabs) pre-equilibrated with buffer B + 15 mM β-ME. The column was then washed with 3 CV of buffer B before being eluted using buffer B + 15mM B-ME and 20mM maltose. The elution was then pooled, spiked with 2 mM EDTA, 5mM dithiothreitol (DTT), and 2mg of recombinant TEV protease before being dialyzed for 16 hours 20 mM HEPES (pH 7.4), 500 mM NaCl, 10mM ϕ3-mercaptoethanol. The dialyzed sample was then applied to a HisTrap Fast Flow IMAC column (GE Healthcare) pre-equilibrated with buffer B. The flowthrough containing the cleaved protein was collected, concentrated using an Amicon concentrator (10K MWCO; Millipore), and applied to a preparative-grade Superdex75 column (GE Healthcare) pre-equilibrated with ITC/Assay Buffer. Monomeric EFPreIQ was then collected and concentrated using an Amicon concentrator (10K MWCO; Millipore) to 1.5 mM for ITC experiments or to 800 μM for kinase assays.

PKAc intended for crystallization was purified using a protocol to separate out the different auto-phosphorylated forms as previously described ((33, 58). Briefly, PKAc was dialyzed against 10 mM potassium-phosphate buffer pH 6.3-6.4 plus 20 mM KCl and 10 mM β-ME before cation exchange. PKAc sample was applied to the SP column equilibrated with 15 mM potassium-phosphate buffer pH 6.3, 20 mM KCl plus 10 mM βME, and subsequently eluted with a gradient of 2% to 28% of buffer containing an additional 1 M KCl over 28 CV. SP column elution peak containing the target protein as confirmed via SDS-PAGE, were concentrated to 1.0 mL and run on a Superdex200 (GE Healthcare) gel filtration column in buffer A (plus 2 mM DTT for PKAc).

### X-Ray Crystallography

The two major SP elution peaks for PKAc were used independently for crystallographic screening **(Fig. S7)**. The two peaks likely correspond to two different species of PKAc: the first eluted species having more autophosphorylation sites than the second. Based on the electron densities we observed for ApoPKAc/AMP-PNP structures (peak 1) versus Complex 2 (peak 2) we presume that the first peak corresponds to a species that is phosphorylated at residues Ser139, Thr197, and Ser338, while the second species is only phosphorylated at residues Thr197 and Ser338.

We obtained crystals in the presence of the Ca_V_1.2 Ser1981 peptide (RGFLRSASLGRRASFHL) using both forms of PKAc (**Table S1**). Peak 1 resulted in the formation of complex 1 (C1) crystal, and peak 2 formed crystal complex 2 (C2). All proteins used for crystallographic screening were concentrated and buffer exchanged with 20 mM bicine pH 8.0, 150 mM ammonium acetate, 4 mM TCEP. The PKAc samples were mixed with synthetic peptide either directly in the lyophilized powder or in a 5mM stock solution in the same buffer. Final PKAc concentration for C1 crystal form was 300 µM with AMP-PNP, MgCl_2_, and Ser1981 peptide at molar ratio of 1:10:17:10. For the peptide co-crystal structure C2 final PKAc concentration was 250 µM in presence of AMP-PNP, MgCl2, and Ser1981 peptide for a molar ratio of 1:10:10:10. We also screened 300 µM PKAc (peak 1) in presence of a shorter Ser1718 peptide DIGPEIRRAISGDL, AMP-PNP, and MgCl_2_ at a molar ratio of 1:10:10:10. No peptide was found to be bound and this thus yielded the apoPKAc crystal form. 96-well plate low volume crystallization plates (Hampton Research) were all set up at room temperature using sitting drop method with ratios 1:1 and 1:2 for precipitant to protein, using a Phoenix crystallization robot (Art Robbins Instruments). All crystal plates were immediately stored at 4°C. For all structures the best diffracting crystals originated from the 1:2 precipitant to protein ratio and were transferred to a drop supplemented with 25% ethylene glycol as cryo-protectant.

Crystals were harvested and frozen in liquid nitrogen using Hampton or MiTeGen MicroMounts cryoloops. All diffraction datasets were processed using HKL2000 (HKL Research Inc.). Best diffracting crystals for Complex1 appeared in condition 64 of ProComplex crystal screen (QIAGEN) with the following formulation: 0.1 M HEPES pH 7.0, and 18% (w/v) PEG 12,000 Da. Crystals for Complex2 appeared in condition 79 of JCSG+ crystal screen (QIAGEN) with the following formulation: 0.1 M Succinic acid pH 7.0, and 15% (w/v) PEG 3,350 Da. The apoPKAc crystal was obtained in condition 64 of Classics crystal screen (QIAGEN) with the following formulation: 0.1 M HEPES pH 7.5, 10% (w/v) PEG 8,000 Da. Datasets for Complex1 and apoPKAc were collected at Stanford Synchrotron Radiation Light source (beamlines 12-2 and BL9-2, respectively) at a wavelength of 0.979Å, using a Dectris Pilatus3 6M detector, while data for the Complex2 crystal form were collected at the Advanced Photon Source (APS, 23 ID D) at a wavelength of 1.033Å, also equipped with a Dectris Pilatus3 6M detector. All structures were solved via molecular replacement in Phaser^61^, using PKAc derived from PDB 6MM5 as a search model. Restrained and translation, libration and screw (TLS) refinement was carried out using PHENIX^62,63^.

### ADP-GloMax^TM^ Kinase Assays

Kinase assays were conducted using an ADP-Glo™ Kinase Assay Kit (Promega). Reagents were prepared as described in the kit, aliquoted, and frozen at −20°C prior to use. Reactions were conducted in 384-well solid white polystyrene microplates (Corning). Peptide substrates and EFPreIQ were dissolved and/or diluted to 800 μM using 50 mM HEPES (pH 7.4), 150 mM KCl, 5 mM MgCl_2_, and 4 mM TCEP. Peptides containing tryptophan had their concentrations determined via absorbance at a wavelength of 280nm using an extinction coefficient of 5500 M^−1^cm^−1^. For peptides lacking tryptophan, the amount of volume needed to achieve a concentration of 800 μM was calculated prior to dissolving. The peptides/substrate were two-fold serially diluted using the same buffer to create eleven reactions per curve ranging from 0 to 256 μM. ATP was added to each reaction to a achieve a final concentration of 1 mM. Reactions were initiated by the addition of thawed PKAc at a final concentration of 3 nM at a final well volume of 20 μL. After allowing reactions to occur at room temperature for 30 minutes, 5 μL from each reaction in a curve was taken and mixed with 5 μL ADP-Glo Reagent. For one experiment, this was done three times (three replicates). After 40 minutes, 10 μL of Kinase Detection Reagent was added to each well and, after waiting 60 minutes, relative luminescence unit (RLU) were measured using a VICTOR X4 Multilabel Plate Reader (Perkin Elmer).

### Kinetic Data Acquisition and Processing

The RLU values were converted to specific activity values via interpolation of a standard curve that was created in accordance with the manufacturer’s protocol. The replicates were averaged and plotted against peptide concentration using GraphPad 9 (Prism) software. The Michaelis-Menten kinetics fitting function was used to generate a non-linear regression to calculate both a K_m_ constant and V_max_ value. Figures displaying data points and Michaelis-Menten curves depict the average specific activity values of the three to four experiments (n=3 or n=4) per peptide (n=4 for Ser1981 and Ser272 peptides, n=3 for all others). Replicates resulting from experimental error were removed from the dataset. In some cases, all three replicates were removed resulting in only two data points for a peptide concentration [Kemptide: 16 μM, Ser1718 Peptide: 2 μM, EFPreIQ (Ser1535): 128 μM, Ser38 Peptide: 2 μM, Ser300 Peptide: 128 μM; 4 μM; and 1 μM] or three data points for the Ser272 peptide (128 μM).

### Isothermal Titration Calorimetry

Isothermal titration calorimetry experiments were conducted using a MicroCal iTC200 (GE Healthcare, now Malvern) instrument. The ITC/Assay buffer was used to dissolve peptides to a concentration of 1.5-4.3 mM. Substrate concentrations were optimized and chosen based on how much was needed to achieve sufficient signal to noise ratio. Rad Ser25 peptide contained a tryptophan residue, which allowed us to determine its concentration via 280nm absorbance readings using an extinction coefficient of 5500 M^−1^cm^−1^. For peptides lacking a tryptophan or tyrosine residue (Ser1981 peptide, Ser1718 peptide, Ser272 peptide, and Ser300 peptide), the amount of volume needed to achieve the desired concentration was calculated prior to diluting. Peptides were received as powder in 1mg aliquots and using molecular weights of the peptide, the number of moles in each aliquot was determined. The number of moles was used to determine how much volume was required to achieve the desired concentration. Adenylyl-imidodiphosphate (AMP-PNP, Millipore Sigma) was dissolved in ITC/Assay buffer to a concentration of 100 mM, aliquoted in 20 μL volumes and stored at −70°C. Prior to each run, both titrant and titrand solutions were supplemented with AMP-PNP to a final concentration of 500 μM. The titration experiments were conducted at 4°C with a stirring speed of 500 RPM. Each titration experiment was comprised of 20 injections with 230 seconds in between each injection. Each injection totalled 2.0 μL over 4s except the first injection being 0.4 μL over 0.8s. Background runs of peptide/substrate into ITC/Assay buffer were conducted to generate a linear regression using the isotherm to subtract background heats from experimental runs. Peptides displaying large background heats were dialyzed in ITC/Assay buffer for 60 minutes to remove contaminants remaining from peptide synthesis. The peptide into PKAc runs (experimental runs) utilized the same settings. The data was processed using Origin (Version 7.0, OriginLab) software. For runs utilizing peptides that did not provide a significant absorbance at 280nm, the concentrations were manually adjusted to achieve a stoichiometry of N=1 (Rad Ser272 and Ca_V_1.2 Ser1981 peptides).

## Data Availability

The atomic coordinates and structure factors for the catalytic subunit of protein kinase A in complex with CaV1.2 S1981 peptide complex1 and 2, and the ApoPKA/AMP-PNP structure have been deposited in the Protein Data Bank with accession codes PDB: 8UKP, 8UKO, and 8UKN, respectively (http://www.rcsb.org/).

### Acknowledgements

We acknowledge Jason Rogalski (UBC) for mass spectrometry, and Dr. Geoffrey Hammond (UBC) for plate reader use. We also thank the support staff at the Advanced Photon Source (Chicago) GM/CA-CAT beamline 23-ID-D, the Stanford Synchrotron Radiation Lightsource (Menlo Park)

## Author Contributions

R.Y, O.H-G., C.X. and C.M. performed experiments. R.Y., O.H-G. and F.V.P. wrote the manuscript. F.V.P. supervised the project.

## Funding and Additional Information

This work is funded by infrastructure funding from the Canadian Fund for Innovation (CFI) and the BC Knowledge Development Fund (BCKDF) to F.V.P. We also acknowledge an operating grant from the Canadian Institutes for Health Research (CIHR) (PJT-153305) to F.V.P. and fellowships from the CIHR and the Michael Smith Foundation for Health Research (MSFHR) to O.H.-G.

