## Supporting Information for "Crystallographic, kinetic, and calorimetric investigation of PKA interactions with L-type calcium channels and Rad GTPase"

```
CaV1.2_Human:(Q13936-1)             DYLTRDWSILGPHHLDEFKRIWAEYDPEAKGRIKHLDVVTLLRRIQPPLGFGKLCPHRVA        1577
CaV1.3_EFPreIQ(S1535):(Q01668-1)    DYLTRDWSILGPHHLDEFKRIWSEYDPEAKGRIKHLDVVTLLRRIQPPLGFGKLCPHRVA        60
                                    **********************:*************************************

CaV1.2_Human:(Q13936-1)             CKRLVSMNMPLNSDGTVMFNATLFALVRTALRIKTEGNLEQANEELRAIIKKIWKRTSMK        1647
CaV1.3_EFPreIQ(S1535):(Q01668-1)    CKRLVAMNMPLNSDGTVMFNATLFALVRTALKIKTEGNLEQANEELRAVIKKIWKKTSMK        120
                                    *****:*************************:****************:******:****

CaV1.2_Human:(Q13936-1)             LLDQVVPPAGD         1658
CaV1.3_EFPreIQ(S1535):(Q01668-1)    LLDQVVPPAGD         131
                                    ***********
```

**Figure S1. EFPreIQ Sequence Comparisons of $Ca_V$1.2 and $Ca_V$1.3.** Partial sequence alignment of residues 1527-1658 of the $Ca_V$1.2 human sequence (uniprot: Q13936-1) to the $Ca_V$1.3 EFPreIQ (S1535) construct based on the human sequence of $Ca_V$1.3 (uniprot: Q01668-1). The Ser1535 residue in human $Ca_V$1.2 is indicated (bold and underlined).

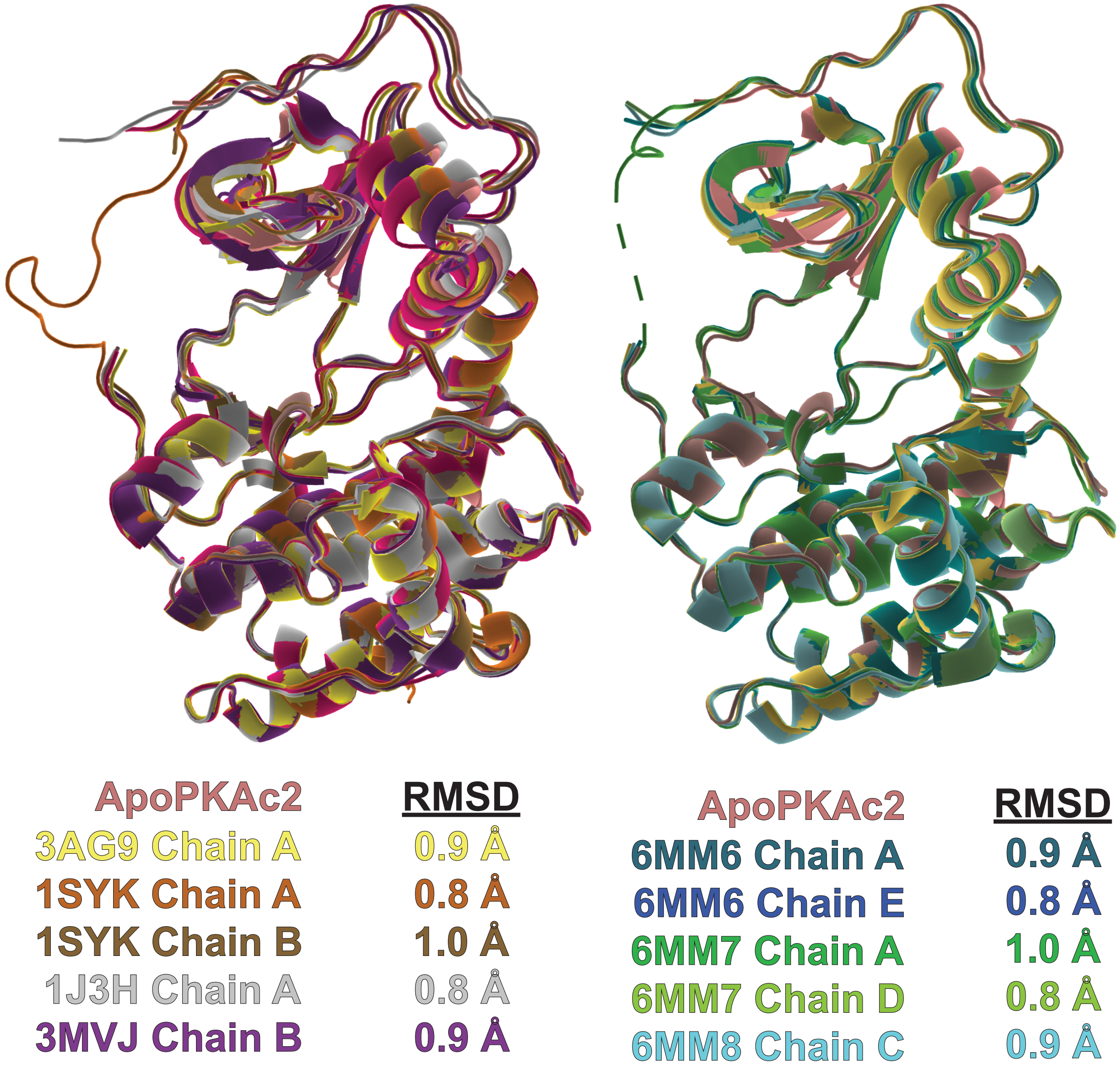


**Figure S2. Structures with the highest structural similarity to ApoPKAc2 molecule (PDB: 8UKN, Chain F).** The top closest structural matches of ApoPKAc2 according to Z-score with RMSD values 1.0 Å or lower overlaid on the ApoPKAc2 structure. The top apo and binary (inhibitor peptide-bound, nucleotide-unbound) structures are shown on the left and the top ternary structures (complexes including AMP-PNP and RyR2 substrate) on the right.

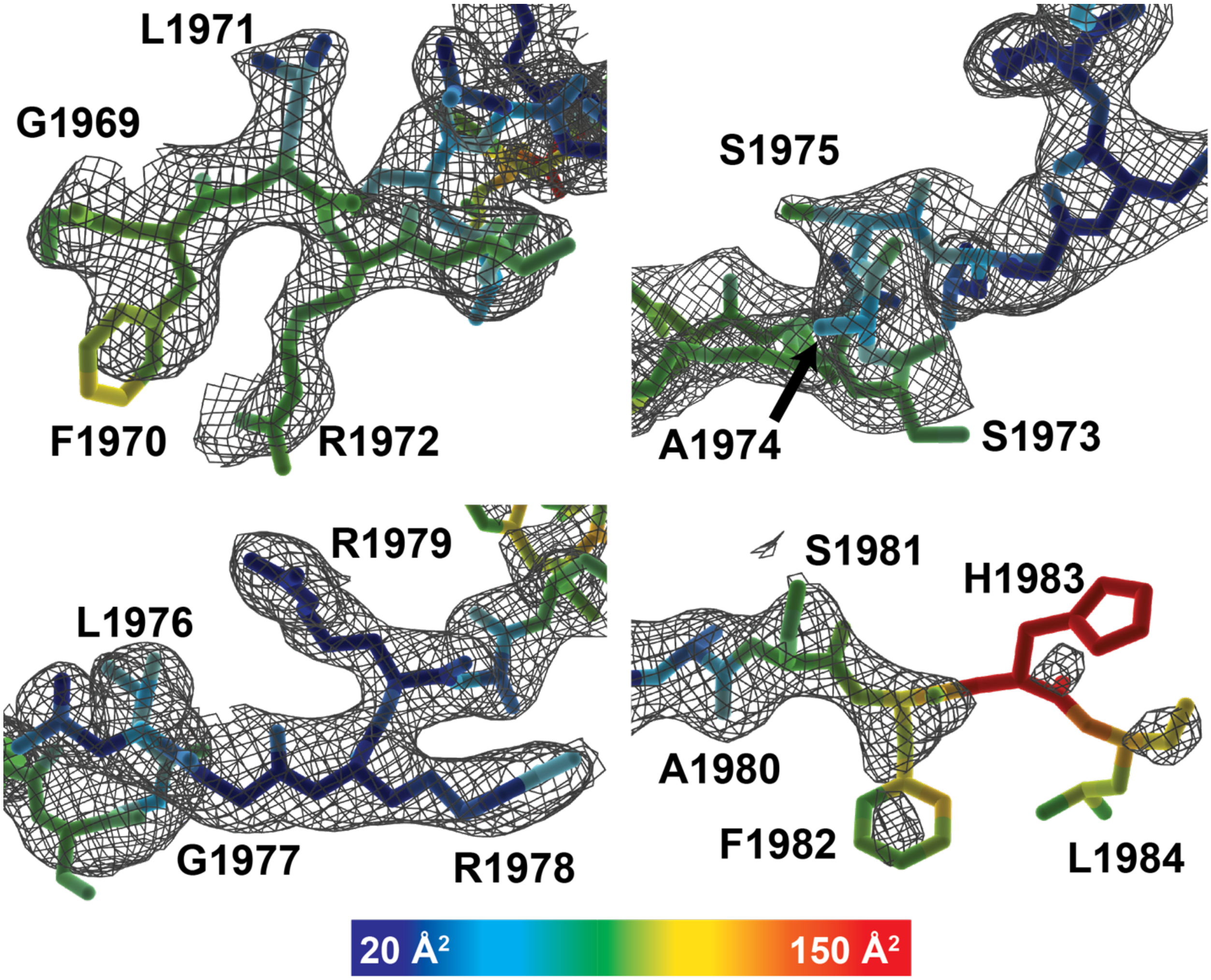


**Figure S3. Density and B-factor colouring of the Ca$_V$1.2 peptide in Complex1**. The 2Fo-Fc density map for the peptide is contoured to 0.7σ. The peptide is coloured according to atom B-factors following the scale shown at the bottom.

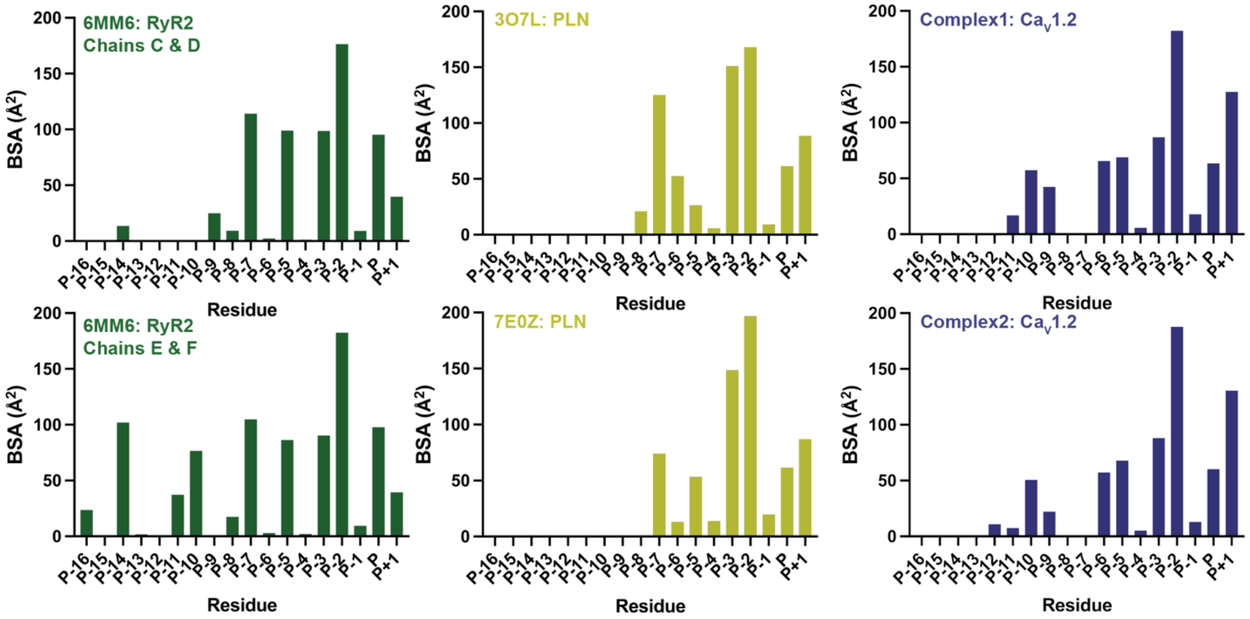


**Figure S4. Buried surface area plots of PKAc:substrate complexes.** Graphs depict how much each residue in each substrate is buried by PKAc.

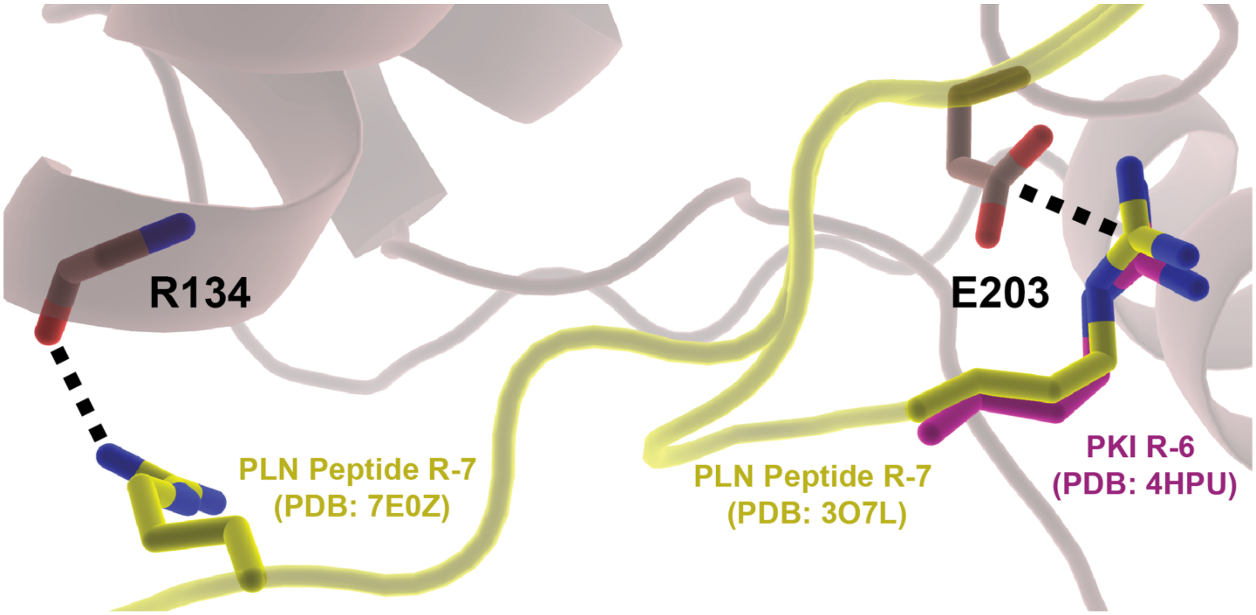


**Figure S5. Differences between R-7 residues of PKAc:PLN ternary structures.** The positions of the R-7 side chains in both PKAc:PLN structures are shown. The R-7 of one PLN structure (PDB: 3O7L) is positioned identically to R-6 of PKI (PDB: 4HPU).

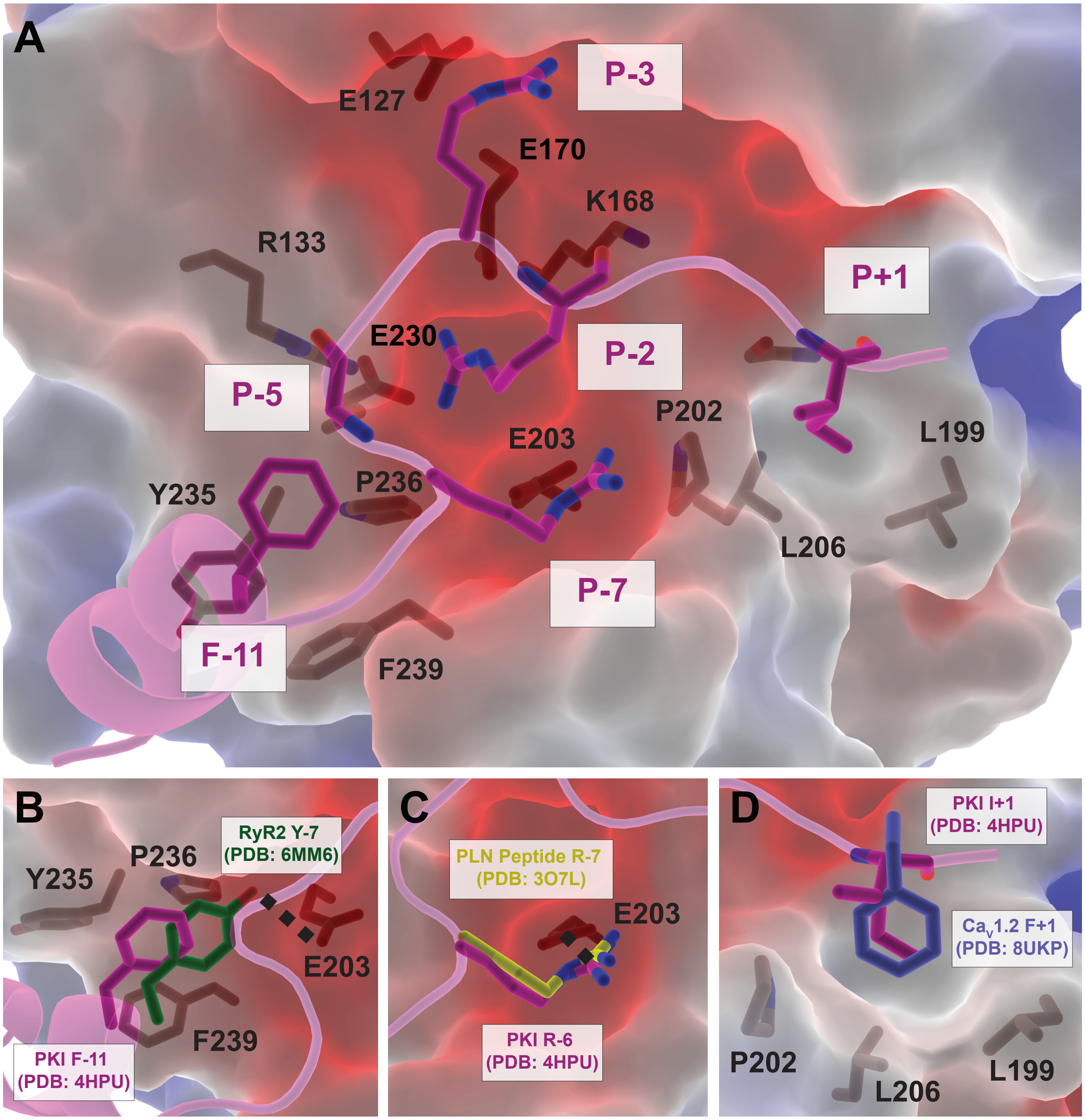


**Figure S6. Interactions of PKI peptide with PKAc active site in comparison with PKAc: substrate complexes.** ***A,*** PDB: 4HPU structure depicts PKAc in complex with substrate competent-PKI. PKAc is coloured according to electrostatic potential. PKI is depicted in cartoon representation. PKAc and PKI residues involved in interactions are shown as sticks. ***B,*** F-11 of PKI slots into a shallow hydrophobic groove in the PKAc active site formed by Y235, P236, and F239. Y-7 of RyR2 (PDB: 6MM6) engages similarly with an additional hydrogen bond to E203. ***C,*** The R-7 in one PKAc:PLN ternary structure (PDB: 3O7L) is positioned nearly identically to R-6 of PKI and forms a salt bridge with E203. ***D,*** A hydrophobic pocket formed by P202, L206, and L199 of PKAc is occupied by I+1 in PKI and F+1 in $Ca_V1.2$ (PDB: 8UKP).

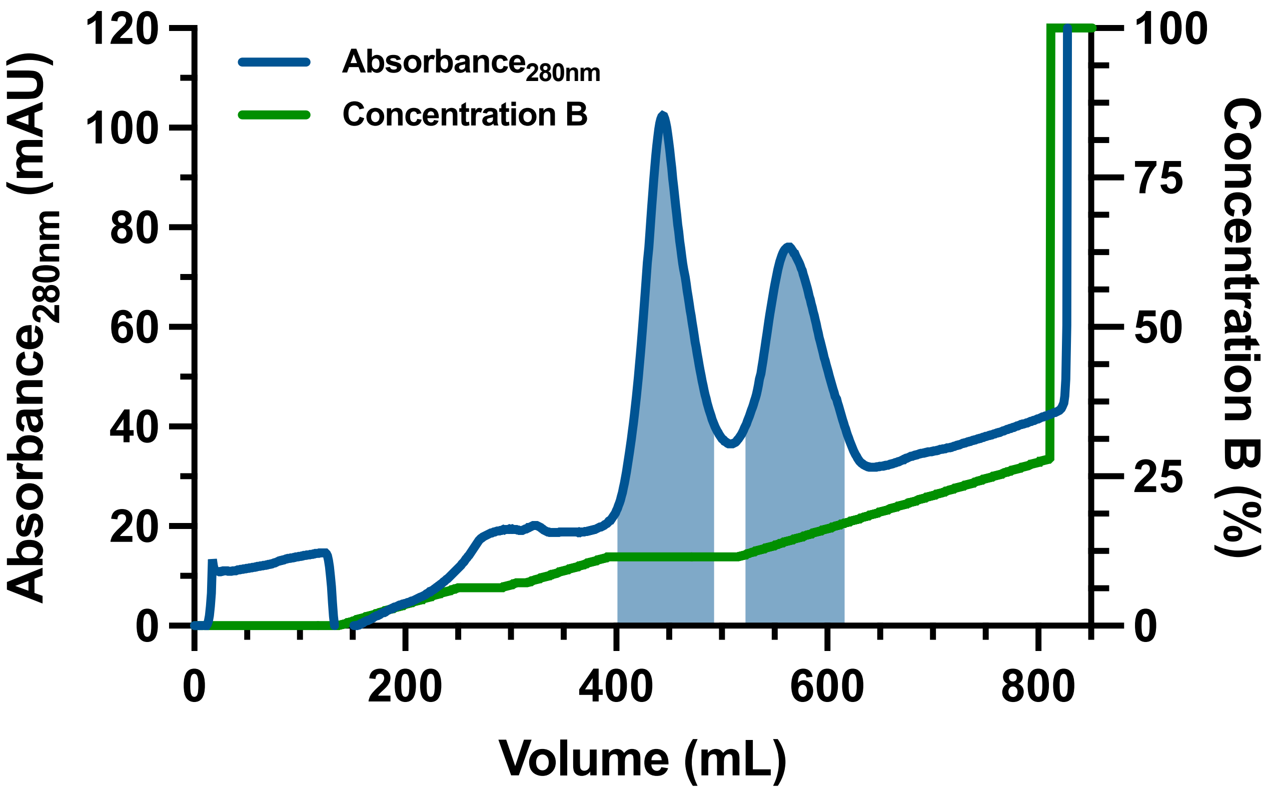


**Figure S7. Cationic Exchange Chromatograph of PKAc.** A representative chromatogram of the chromatography step is shown with the $UV_{280}$ (mAU) shown in blue (left y-axis) and percent concentration of buffer B (15 mM potassium-phosphate buffer [pH 6.3], 1 M KCl, and 10 mM βME) applied to the column shown in green (right y-axis). Two predominant species elute closely together, requiring the holding of the salt gradient to ensure sufficient separation of the two species.

**Table S1. Crystallographic Table.**

| **Crystal** | **PKAc-S1981 Complex1** | **PKAc-S1981 Complex2** | **Apo/AMP-PNP PKAc** |
|---|---|---|---|
| PDB codes | 8UKP | 8UKO | 8UKN |
| Resolution (Å) | 35.0–2.85 (2.92–2.85)[a] | 35.0–2.99 (3.06–2.89)[a] | 35.0–2.75 (2.81–2.75)[a] |
| Space group | $P\ 4_1 2_1 2$ | $I\ 4_1$ | $P2_1$ |
| *a* (Å) | 118.63 | 118.058 | 61.8 |
| *b* (Å) | 118.63 | 118.058 | 139.8 |
| *c* (Å) | 57.11 | 58.0 | 107.4 |
| α, β, γ (°) | 90.0, 90.0, 90.0 | 90.0, 90.0, 90.0 | 90.0, 102.8, 90.0 |
| Volume $Å^3$ | $8.05 \times 10^5$ | $8.08 \times 10^5$ | $9.05 \times 10^5$ |
| Wavelength (Å) | 0.979 | 0.979 | 1.033 |
| No. of molecules/A.U. | 1 | 1 | 4 |
| Unique reflections | 9,956 | 9,121 | 44,968 |
| Redundancy | 13.6 (14.1) | 4.7 (4.6) | 3.7 (3.3) |
| 〈I/σ(I)〉 | 35.4 (1.37) | 25.2 (1.4) | 16.1 (2.15) |
| CC1/2 (%)[b] | 99.7 (57.1) | 99.5 (34.2) | 99.7 (83.0) |
| *R*pim (%)[c] | 2.80 (100.9) | 3.00 (58.3) | 4.50 (29.7) |
| Completeness (%) | 100.0 (100.0) | 99.9 (100.0) | 97.2 (85.1) |
| Protein atoms | 2,678 | 2,781 | 10,587 |
| Water oxygen atoms | 12 | 8 | 136 |
| Ligands or ions | 33 | 33 | 35 |
| Ramachandran outliers | 0 | 0 | 0 |
| Solvent content (%)[d] | 53.1 | 51.5 | 58.8 |
| Wilson *B*-factor ($Å^2$) | 52.8 | 50.9 | 42.7 |
| ***Refinement*** | | | |
| $R_{work}$ (%) | 24.0 | 21.0 | 19.5 |
| $R_{free}$ (%)[e] | 28.1 | 25.9 | 23.2 |
| RMSD bond lengths (Å) | 0.002 | 0.002 | 0.004 |
| RMSD bond angles (°) | 0.48 | 0.43 | 0.63 |
| Mean *B*-factor ($Å^2$) | 65.0 | 64.0 | 47.0 |
| Protein atoms ($Å^2$) | 65.7 | 64.7 | 47.1 |
| Water atoms ($Å^2$) | 33.8 | 28.5 | 35.8 |
| Ligands and ions ($Å^2$) | 70.9 | 41.5 | 91.7 |

*Two protein kinase A catalytic domain (PKAc) crystal structures were solved in complex with a peptide containing the CaV1.2 S1981 target site. These are referred to as crystal form 1 and 2 (CF1 and CF2). The peptide used was RGFLRSASLGRRASFHL (1968-1984) respectively. RMSD, root mean square deviation. NA: Not applicable.*

[a] *Values in the parentheses refer to the highest resolution shell*

[b] $CC_{1/2} = \sigma_i^{\ 2} / (\sigma_i^{\ 2} + \sigma_j^{\ 2})$*, where* $\sigma_j$ *denotes mean error for half-dataset* [3]

[c] $Rpim = \sum_{hkl} (1/n - 1)^{1/2} \sum_i |I_{hkl,\ i} - [I_{hkl}]| / \sum_{hkl} \sum_i I_{hkl,\ I}$*, where* $[I_{hkl}]$ *is the average of Friedel-related observations (i) of a unique reflection (hkl)*

[d] *Estimated using sfcheck* [4]

[e] *5% of reflections were omitted for* $R_{free}$ *calculations*

**Table S2. Distances Between Residues Used to Determine Open/Intermediate/Closed States**

| Crystal Structure | Molecule Name (Chain) | G52-D166 (Cα-Cα) | S53-G186 (Cα-Cα) | H87−T197 (Nε2-PO4) | E170−Y330 (C=O−OH) | State |
|---|---|---|---|---|---|---|
| Apo/AMP-PNP PKAc (PDB: 8UKN) | ApoPKAc 1 (C) | 19.5 | 14.1 | 7.3 | n/a | Open |
| | ApoPKAc 2 (F) | 19.1 | 14.0 | 6.6 | n/a | Open |
| | ApoPKAc 3 (H) | 19.6 | 14.2 | 8.0 | n/a | Open |
| | AMP-PNP PKAc (D) | 17.4 | 13.1 | 6.6 | n/a | Intermediate |
| PKAc-S1981 Complex1 (PDB: 8UKP) | Complex 1 (E) | 16.1 | 12.9 | 3.1 | n/a | Intermediate |
| PKAc-S1981 Complex2 (PDB: 8UKO) | Complex 2 (E) | 16.5 | 13.5 | 5.5 | 9.1 | Intermediate |

*Measurements done in PyMol.*

**Table S3. Interactions Between the $Ca_V1.2$ S1981 Peptide and PKAc**

| $Ca_V1.2$ S1981 Peptide Residue | Interaction Type | Complex 1 PKAc Residues (PDB: 8UKP) | Complex 2 (PDB: 8UKO) |
|---|---|---|---|
| G1969 | VDW | F239 | - |
| F1970 | VDW | F239 | F239 |
| L1971 | VDW | Y235, F239 | Y235, F239 |
| R1972 | SB | - | D241 |
| | $HB^{NH1}$ | - | $D241^{OD2}$ |
| | VDW | D241 | D241 |
| S1975 | VDW | R133, E203, P236, F239 | R133, E203, P236, F239, A240 |
| L1976 | $HB^{O}$ | $R133^{NE}$ | $R133^{NE}$ |
| | VDW | F129, R133, G234, Y235, P236, | F129, R133, G234, Y235, P236 |
| G1977 | VDW | F129, E170 | F129, E170, R133 |
| R1978 | SB | E127 | E127 |
| | $HB^{NE}$ | $E127^{OE2}$ | $E127^{OE1}$ |
| | $HB^{NH2}$ | $E127^{OE2}$ | $E127^{OE2}$ |
| | VDW | E127, F129, E170, N171 | E127, F129, E170 |
| R1979 | SB | E170, E230 | E170, E203, E230 |
| | $HB^{N}$ | $E170^{OE2}$ | $E170^{OE2}$ |
| | $HB^{NE}$ | $E170^{OE2}$ | $E170^{OE2}$ |
| | $HB^{NH1}$ | - | $E203^{OE1}$ |
| | | - | $E230^{OE1}$ |
| | | $E230^{OE2}$ | - |
| | $HB^{NH2}$ | $E170^{OE1}$ | $E170^{OE2}$ |
| | | $E230^{OE2}$ | $E230^{OE2}$ |
| | $HB^{O}$ | $K168^{NZ}$ | $K168^{NZ}$ |
| | VDW | F129, R133, K168, P169, E170, T201, E203, Y204, E230, P236 | F129, R133, K168, P169, E170, T201, E203, Y204, E230, P236 |
| A1980 | VDW | K168, G200, T201 | K168, G200, T201, P202 |
| S1981 | VDW | D166, K168, F187, G200, T201 | D166, K168, F187, G200, T201 |
| F1982 | $HB^{N}$ | $G200^{O}$ | $G200^{O}$ |
| | $HB^{O}$ | $G200^{N}$ | - |
| | VDW | F187, L198, C199, G200, T201, P202, L205, Y247 | F187, L198, C199, G200, T201, P202, L205, Y247 |

Hydrogen bonds and salt bridge interactions were determined using PDBePISA[10]. Van der Waals (VDW) Interactions were determined using CONTACT[11]. Hydrogen bonding donor and acceptor atoms are listed for hydrogen bonds as superscripts. The hydrogen bond (HB) distance cut-off was set to 3.5 Å and the salt bridge (SB) distance cut-off (measured from points at each residue's charge centre) was set to 4.0 Å. Interactions unique to each complex are listed in red.

**Table S4. Human $Ca_V1.2$ sequence variants of PKAc binding region.**

| Mutation | Disease | Source | Predicted Effect |
| --- | --- | --- | --- |
| G1969A | Long QT syndrome | Clinvar/gnomAD | Affects vdW contacts with F239 of PKAc |
| R1972H | Long QT syndrome | Clinvar/gnomAD | Ablates vdW contacts with D241 side chain of PKAc |
| R1972C | Timothy syndrome, Cardiovascular phenotype (not provided) Long QT syndrome, Arrhythmogenic right ventricular cardiomyopathy, History of neurodevelopmental disorder (not specified) | Clinvar/gnomAD | Ablates vdW contacts with D241 of PKAc |
| A1974P | Cardiovascular phenotype (not provided), Long QT syndrome, Timothy syndrome, Brugada syndrome | Clinvar/gnomAD | Affects the positioning of $Ca_V1.2$ by restricting backbone movement, and perturbing binding. |
| G1977S | Timothy syndrome, Long QT syndrome, Brugada syndrome | Clinvar | Introduce steric impediments, abrogating vdw contacts with F129 and E170. |
| G1977D | Long QT syndrome | Clinvar/gnomAD | Introduce steric impediments, abrogating vdw contacts with F129 and E170. |
| R1978Q | Timothy syndrome, Brugada syndrome, Cardiovascular phenotype, Long QT syndrome | Clinvar/gnomAD | Affects salt bridging interaction of R1978 with electronegative pocket of PKAc (E170, E203, E230). |
| R1979K | Long QT syndrome | Clinvar | Affects salt bridge formed with E127 of PKAc. |
| S1981P | Long QT syndrome | Clinvar | Phosphorylation-incompetent. |
| S1981F | Long QT syndrome | Clinvar/gnomAD | Phosphorylation-incompetent. |

Sequence variants in the Ser1981 peptide according to the Clinvar and gnomAD databases, showing the disease description and prediction of the impact of the variants based on the structure presented in this manuscript.
